# Multimodal single-cell analyses reveal molecular markers of neuronal senescence in human drug-resistant epilepsy

**DOI:** 10.1101/2024.12.07.627313

**Authors:** Qianqian Ge, Jiachao Yang, Fei Huang, Xinyue Dai, Chao Chen, Jinxin Guo, Mi Wang, Mengyue Zhu, Yijie Shao, Yuxian Xia, Yu Zhou, Jieqiao Peng, Suixin Deng, Jiachen Shi, Yiqi Hu, Huiying Zhang, Yi Wang, Xiaoqun Wang, Xiao-Ming Li, Zhong Chen, Yousheng Shu, Jun-Ming Zhu, Jianmin Zhang, Ying Shen, Shumin Duan, Shengjin Xu, Li Shen, Jiadong Chen

**Author notes:** Lead contact. Jiadong Chen. These authors contributed equally to this work (Qianqian Ge, Jiachao Yang, Fei Huang, Xinyue Dai, Chao Chen). Corresponding authors: Jiadong Chen, Li Shen, Shengjin Xu; Address correspondence to: Jiadong Chen, Department of Neurobiology, Zhejiang University School of Medicine, 866 Yuhangtang Road, Hangzhou 310058, China. Or to: Li Shen Zhejiang University, 866 Yuhangtang Road, Na Mi Bldg 425, Hangzhou 310058, China. Or to: Shengjin Xu, Institute of Neuroscience, Center for Excellence in Brain Science and Intelligence Technology, Chinese Academy of Sciences, 320 Yueyang Road, Shanghai 200031, China.

## Abstract

The histopathological neurons in the brain tissue of drug-resistant epilepsy exhibit aberrant cytoarchitecture and imbalanced synaptic circuit function. However, the gene expression changes of these neurons remain unknown, making it difficult for the diagnosis or to dissect the mechanism of drug-resistant epilepsy. By integrating whole-cell patch clamp recording and single-cell RNA sequencing approaches, we identified a transcriptionally distinct subset of cortical pyramidal neurons. These neurons highly expressed genes *CDKN1A* (P21), *CCL2* and *NFKBIA* that associated with mTOR pathway, inflammatory response and cellular senescence. We confirmed the expression of senescent marker genes in a subpopulation of cortical pyramidal neurons with enlarged soma size in the brain tissue of drug-resistant epilepsy. We further revealed the expression of senescent cell markers P21, P53, COX2, γ-H2AX, β-Gal and reduction of nuclear integrity marker Lamin B1 in histopathological neurons in the brain tissue of drug-resistant epilepsy patients with different pathologies, but not in control brain tissue with no history of epilepsy. Additionally, chronic, but not acute, epileptic seizures induced senescent markers expression in cortical neurons in mouse models of drug-resistant epilepsy. These results provide important molecular markers for histopathological neurons and new insights into the pathophysiological mechanisms of drug-resistant epilepsy.

## Introduction

Medically intractable epilepsy patients respond poorly to currently available anti-epileptic drugs (1). Histopathological examinations of brain tissue surgically removed from the epileptic focus revealed various anatomical signatures of neuropathologies (2, 3). For example, dysmorphic neurons in the grey matter and balloon cells in the white matter were pathological characteristics in drug-resistant epilepsy patients with focal cortical dysplasia (FCD) (4, 5), while neuronal loss and hippocampal sclerosis were characteristic in the epileptic focus of drug-resistant epilepsy patients with temporal lobe epilepsy (TLE) (2, 3, 6). Understanding the gene expression changes in these cortical pyramidal neurons will be important both for the diagnosis and dissecting the mechanism of seizure genesis in these patients (2, 7). Bulk gene expression studies of the brain tissue in the epileptic focus of drug-resistant epilepsy using microarray or RNA sequencing showed upregulation of genes associated with neuroinflammation, restructuring of neuronal networks (8–11) and activation of mTOR pathway (12). However, representing a rare cell population within the cortex, the histopathological neurons are difficult to capture or enrich by conventional single-cell genomic analysis following tissue dissociation, thus making it difficult to diagnose or dissect mechanisms of seizure genesis in drug-resistant epilepsy.

Single-cell RNA sequencing has revealed cellular heterogeneity and identified molecular markers of various cell types in the adult brain (13–15), including not only molecular markers of canonical cell types previously defined by anatomical and morphological features, but also transcriptomic cell subtypes in different brain regions and across various species. In addition, single-nucleus RNA sequencing identified disease-associated gene expression changes in specific cell types that were vulnerable in various brain diseases including autism, epilepsy, and multiple sclerosis in the human brain (14, 16, 17). The molecular state of cortical upper layer excitatory neurons was selectively affected in patients with autism and epilepsy (14), but the anatomical, physiological and pathological characteristics of the transcriptomic cell subtypes were yet to be explored. Multi-modal patch-seq analysis integrates the morphological, electrophysiological, and single-cell RNA sequencing approaches to functionally characterize transcriptomic cell subtypes (18, 19). Patch-seq analyses have revealed association between cell-type specific electrophysiological characteristics and genes expression profiles under physiological and disease state (20, 21), but whether this approach could reveal molecular characteristics of histopathological neurons in the brain remains unexplored.

In the present study, we performed patch-seq analysis to investigate the electrophysiological and molecular characteristics of histopathological neurons in brain tissues from the epileptic focus of drug-resistant epilepsy patients. Our results revealed a transcriptomic pyramidal neuron cluster that are associated with the pathological state of cortical neurons in epilepsy. We further validated the expression of senescent marker genes and increase soma volume in sub-population of cortical pyramidal neurons in the brain tissue of drug-resistant epilepsy. These results revealed molecular markers and the senescent state of histopathological neurons in people with drug-resistant epilepsy.

## Results

### Patch-seq analyses of pathological neurons in brain tissue from people with drug-resistant epilepsy

To delineate the electrophysiological and molecular characteristics of histopathological neurons, we integrated whole-cell patch clamp recording and single-cell RNA sequencing analysis (patch-seq) in acute brain slices from postsurgical brain tissues of people with drug-resistant epilepsy. Cortical pyramidal neurons and interneurons are identified under differential interference contrast (DIC) imaging by their intrinsic membrane characteristics, action potential firing pattern and morphological characteristics (Figure 1, A and B). After whole-cell recordings, the cytosol of recorded neurons was aspirated by the recording pipette and subsequently subjected to whole transcriptomic single-cell RNA sequencing (Figure 1A). We collected high quality data from 197 neurons that passed the quality assessment (see *Method* section) and clustered them into four clusters as shown by Uniform Manifold Approximation and Projection (UMAP) analysis (Figure 1C). Cluster 0 neurons highly express markers of cortical interneurons (*GAD1, GAD2, ERBB4, DLX1, LHX6*) and are annotated as interneurons (INT) (Figure 1, C and D). Cluster 1, cluster 2 and cluster 3 neurons highly express cortical pyramidal neuronal markers (*SATB2, CUX2, SLC17A6, SLC17A7, TBR1*) and are annotated as pyramidal neurons (PY1, PY2, PY3) (Figure 1, C, D, and F). We found that genes of the mTOR pathway (*RPS6, RHEB, EIF4E*) are highly expressed in the PY2 pyramidal neurons (Figure 1, D and F), consistent with the activation of mTOR pathway in histopathological neurons in the epileptic focus of drug-resistant epilepsy (5, 12). The immediate early genes including (*EGR2, EGR3, JUN*), dual-specificity kinase / phosphotase MKP-3 (*DUSP6*) are highly expressed in the PY2 pyramidal neurons (Figure 1D and Supplemental Figure 1B), consistent with the expression of activity-dependent genes in cortical neurons in drug-resistant epilepsy (22).

**Figure 1.**
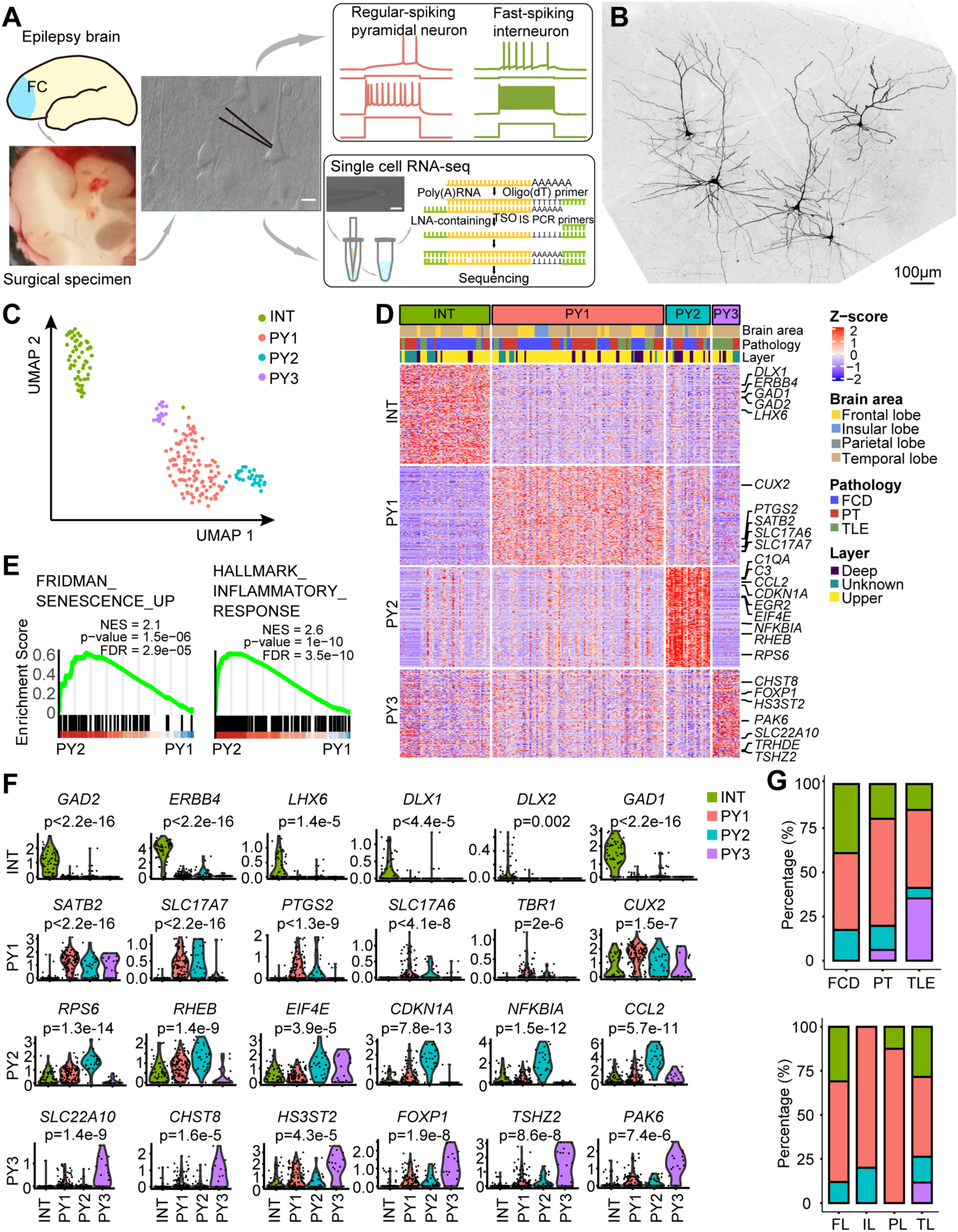
Patch-seq analyses of the molecular and electrophysiological characteristics of cortical neurons in human drug-resistant epilepsy. **(A)** Schematic diagram of patch-seq procedure. Whole-cell patch clamp recording of single neurons were established in acute brain slices from postsurgical brain tissue from people with drug-resistant epilepsy. Example action potential firing traces showed representative regular spiking pyramidal neuron and fast-spiking interneuron. After recordings, the patched neuron was aspirated for subsequent whole-transcriptome RNA sequencing. Scale bar, 10 μm. **(B)** Example image showing the morphology of patch-seq pyramidal neurons by intracellular labeling with biocytin and subsequently immunostained with streptavidin 549. Scale bar 100 μm. **(C)** Distribution of four transcriptomic patch-seq neuron clusters in UMAP plot. INT: interneuron subclass; PY 1 - 3: pyramidal neuron clusters 1 - 3. n = 197 neurons from 36 brain samples. **(D)** Heatmap of top 100 differentially expressed genes (DEGs) of four transcriptomic clusters. **(E)** GSEA analysis of PY2 cluster DEGs showed genes enrichment for cellular senescence and inflammatory response in comparison with PY1 cluster. **(F)** Violin plots showed representative DEGs in INT and PY 1 - 3 neuron clusters. Y axis indicated relative expression level. Kruskal-Wallis test with Dunn’s multiple comparisons test. **(G)** Stacked bar graphs showing the distribution of four transcriptomic neuron clusters in different brain regions (top) and pathologies of drug-resistant epilepsy (bottom), respectively. FCD, focal cortical dysplasia; PT, peri-tumor cortical tissue; TLE, temporal lobe epilepsy. FL, frontal lobe; IL, insular lobe; PL, parietal lobe; TL, temporal lobe.

The gene ontology (GO) analysis for the marker genes of each cluster showed genes highly expressed in the INT interneurons are enriched for telencephalon development and forebrain neurons differentiation (Supplemental Figure 1A). The highly expressed genes in PY1 pyramidal neurons are enriched for synaptic organization and modulation of chemical synaptic transmission, PY2 genes are enriched for translational initiation, positive regulation of cytokine production and cellular response to oxygen levels, and PY3 genes are enriched for intracellular signal transduction and protein phosphorylation (Supplemental Figure 1A). Gene set enrichment analysis (GSEA) further revealed that PY2 pyramidal neurons showed substantial up-regulation of genes related to senescence and inflammatory responses in comparison with PY1 pyramidal neurons (Figure 1E). Cellular senescence is a state of permanent cell cycle arrest and is characterized by upregulation of genes associated with cyclin-dependent kinase inhibitors, DNA damage response, oxidative stress, inflammatory response and senescence-associated secretory phenotype (SASP) factors (23, 24). We found that genes closely related to cellular senescence and inflammation pathway (*CDKN1A, NFKBIA*) and genes encoding the major categories of SASP factors are enriched in the PY2 pyramidal neurons (11, 25, 26). Indeed, we found enrichment of genes related to soluble signaling factors such as interleukins (*IL1A, IL1B*), chemokines (*CCL4, CCL2, CXCL8, CXCL12*), secreted proteases and regulators (*MMP2, CTSB*), and soluble ligands or shed receptors (*ICAM1, TNFRSF1B, TNFRSF12A*) in the PY2 cluster (Figure 1, D and F, and Supplemental Figure 1B). These results suggest that PY2 pyramidal neuron cluster are tightly associated with the pathological state of cortical pyramidal neurons in drug-resistant epilepsy.

### Multiplex Expansion-ASsisted Iterative Fluorescence In Situ Hybridization (EASI-FISH) reveals molecular and physiological profiles of PY1 and PY2 neurons in the brain tissue

Next we examined whether the four transcriptomic neuron clusters exhibited distinct electrophysiological characteristics by quantifying their intrinsic membrane properties and key features of their action potential firing pattern. We extracted 13 electrophysiological parameters measured from intrinsic membrane characteristics and kinetics of action potentials induced by rheobase and suprathreshold current injections (Figure 2, A and B, and *Method*s) (20, 27, 28). We found the INT interneurons exhibited a typical fast spiking action potential (AP) firing pattern (higher AP firing frequency), larger afterhyperpolarization of AP, smaller membrane capacitance (Cm), and higher input resistance (Rin) (Figure 2, A and C), that are characteristic features of cortical interneurons (29). The pyramidal neuron clusters (PY 1-3) exhibited regular-spiking AP firing patterns that are characteristic of cortical pyramidal neurons (Figure 2A) (30). We did not identify significant differences of intrinsic membrane properties or electrophysiological characteristics between the PY1 and PY2 pyramidal neuron clusters (Figure 2, A and C).

**Figure 2.**
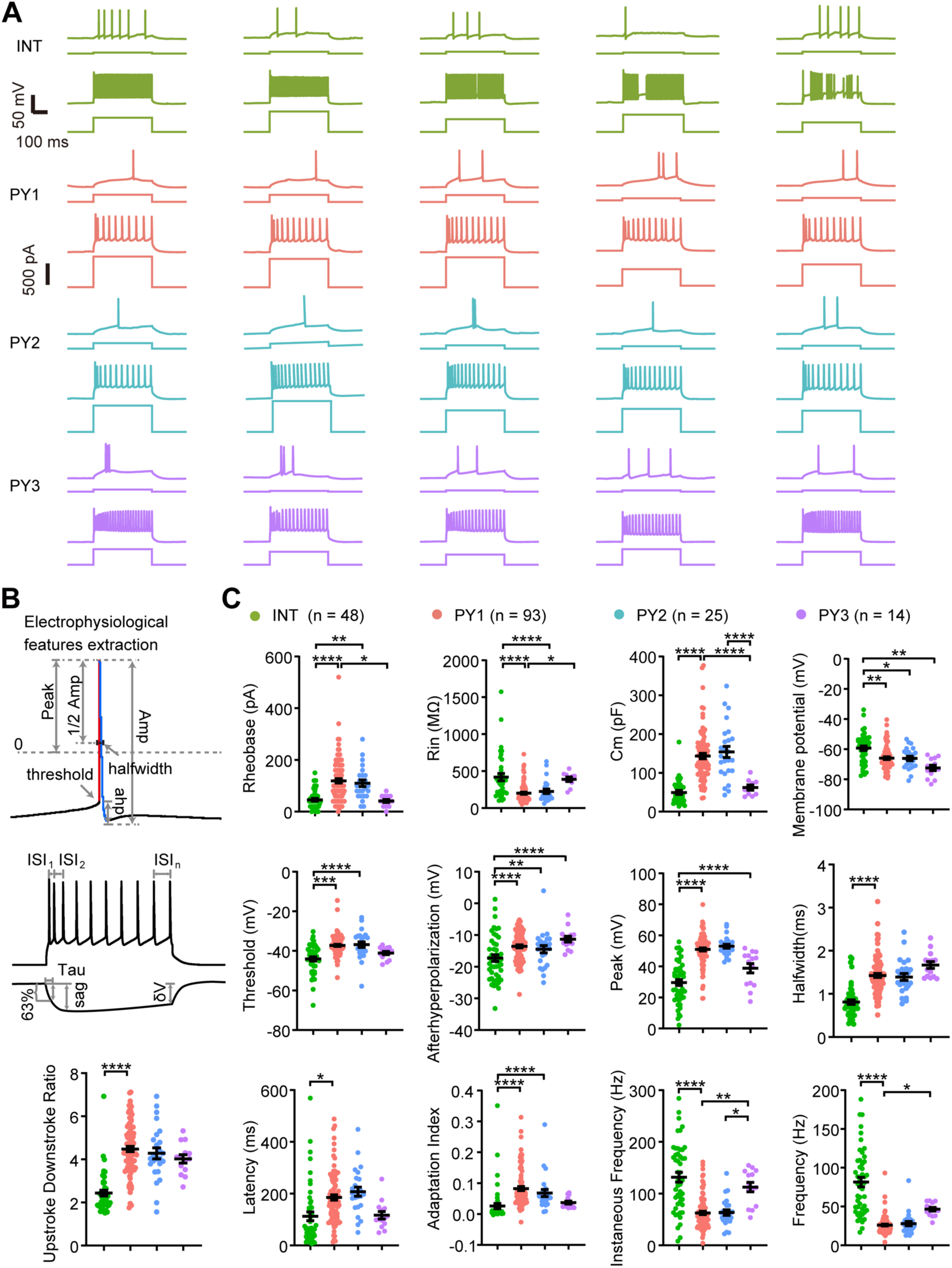
Electrophysiological characteristics of different transcriptomic neuron clusters. **(A)** Representative examples of action potential (AP) firing patterns of four transcriptomic neuron subclasses. Representative AP firing pattern upon rheobase (upper) and suprathreshold (lower) current injection from 5 neurons at each transcriptomic cluster (INT, PY1 - 3). Scale bars: 50 mV, 100 ms in AP firing patterns trace; 500 pA, in stimulation trace. **(B)** Schematic showing extraction of electrophysiological properties of rheobase action potential firing pattern, maximum action potential firing pattern and subthreshold stimulation for quantification analysis (see Methods). Amp, amplitude; Ahp, afterhyperpolarization; ISI, inter-spike interval. **(C)** Statistical analysis of intrinsic membrane characteristics and electrophysiological parameters of four transcriptomic neuron clusters. Rin, input resistance; Cm, membrane capacitance. INT, n = 48; PY1, n = 93; PY2, n = 25; PY3, n = 14. \**P* < 0.05; \*\**P* < 0.01; \*\*\**P* < 0.001; \*\*\*\**P* < 0.0001, Welch’s ANOVA and Tukey multiple comparison test.

To link the morphological and electrophysiological profiles with marker genes expression of PY1 and PY2 neurons in the brain tissue, we integrated patch-clamp recording with multiplex Expansion-ASsisted Iterative Fluorescence In Situ Hybridization (EASI-FISH), which is compatible with the thick brain slices after electrophysiological recording (Figure 3A). We performed whole-cell electrophysiological recording and sparsely loaded individual cortical pyramidal neurons with biocytin in brain tissue from people with drug-resistant epilepsy. Subsequently we reconstructed the morphology of recorded neurons and performed EASI-FISH staining of PY2 marker genes including *CDKN1A*, *NFKBIA*, *CCL2* and PY1 marker gene *CUX2*, as well as *NEFM* and *NEFH* genes that encode neurofilament medium chain and heavy chain, respectively (Figure 3, A and B). We quantified the expression of marker genes and re-cluster the cortical pyramidal neurons solely based on gene expression profiles and identified two clusters of cortical neurons (Figure 3, C-E). After tissue expansion, DAPI staining labeled the full soma of neurons in the brain tissue and we were thus able to quantify the soma size of each individual neurons (31). The cluster 2 neurons had substantially larger soma sizes and increased expression of *CDKN1A, NFKBIA, CCL2, CUX2, NEFM* and *NEFH* compared with cluster 1 neurons (Figure 3, C-I, and Supplemental Figure 3). These results suggest that cluster 2 neurons resemble the molecular features of PY2 neurons and were thus annotated as the molecularly-defined PY2 neurons. In contrast, cluster 1 neurons with smaller soma sizes expressed PY1 marker genes *CUX2* and were annotated as the molecularly-defined PY1 neurons.

**Figure 3.**
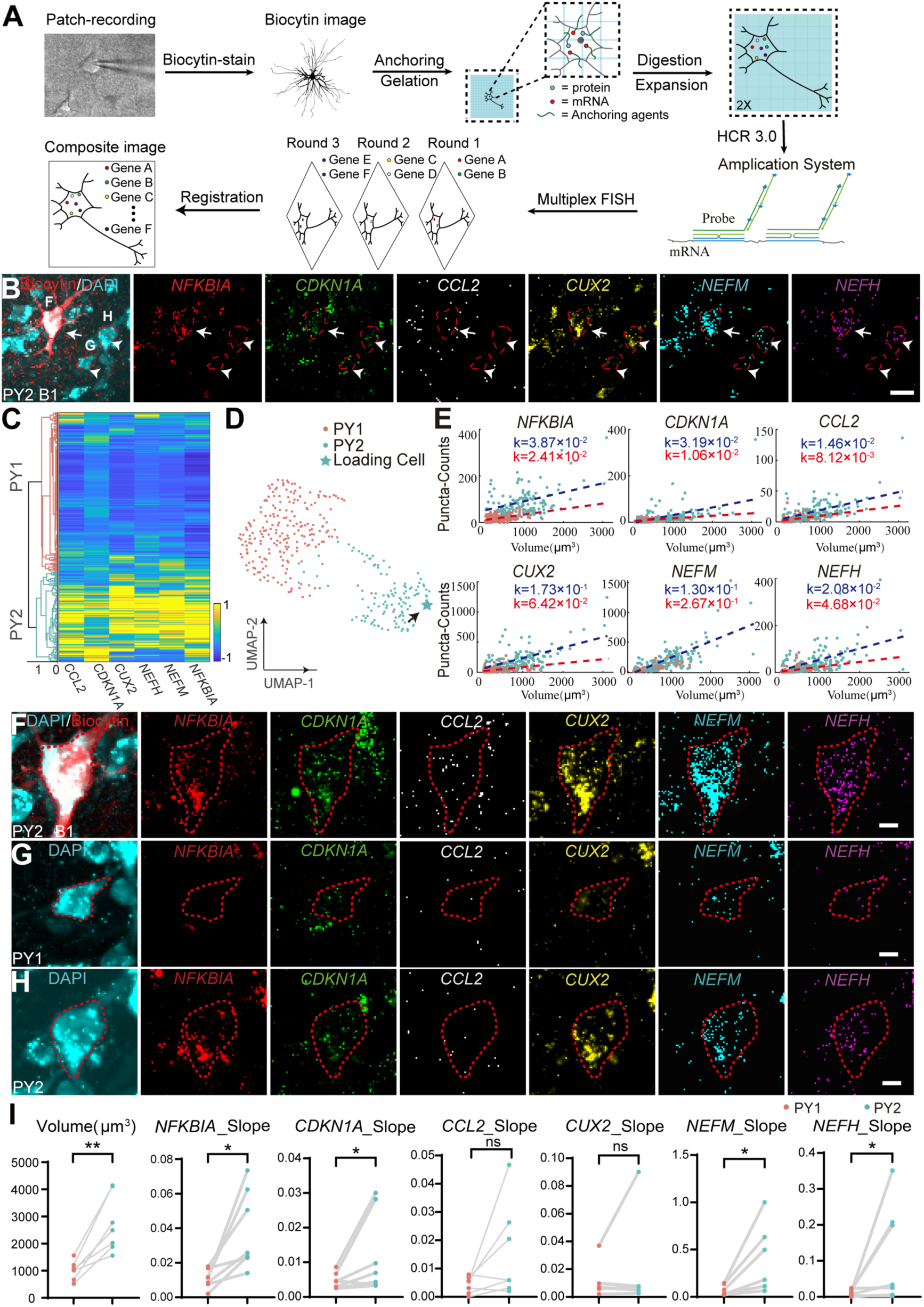
Multiplex EASI-FISH reveals molecularly-defined PY1 and PY2 neurons as well as their electrophysiological and morphological characteristics. **(A)** Schematic diagram illustrating the experimental procedures for loading the morphology of recorded neurons and subsequent staining with multiplex EASI-FISH. **(B)** Representative FISH images showing the expression of six differentially expressed genes (DEGs) *NFKBIA, CDKN1A, CCL2, CUX2, NEFM, NEFH* in a ROI with biocytin loaded PY2 neuron in a brain slice from focal cortical dysplasia. Scale bar, 50 μm. **(C)** Heatmap of six DEGs and Uniform Manifold Approximation and Projection (UMAP) analysis (D) of molecularly-defined PY1 (217 neurons) and PY2 clusters (122 neurons) in an example brain slice. (**E)** Scatter plots showing the gene expression density of six DEGs in PY1 and PY2 neuron clusters. Red dashed line, PY1 cluster density fitting line; Blue dashed line, PY2 cluster density fitting line. **(F-H)** Representative images showing the expression of candidate marker genes in biocytin-labeled PY2 neuron (F), molecularly-defined PY1(G) and PY2(H) pyramidal neurons. **(I)** Statistical results showing the cell size (left) and gene expression density (Slope of density fitting line) (right) in PY1 and PY2 neurons (n = 7688 neurons, 7 brain slices from 4 individual samples). Volume, *P* = 0.0020; *NFKBIA*, *P* = 0.0319; *CDKNIA*, *P* = 0.0469; *CCL2*, *P* = 0.0974; *CUX2*, *P* = 0.5781; *NEFM*, *P* = 0.0361; *NEFH*, *P* = 0.0394. Each data point indicates the mean value from a sample; gray lines connect data points from the same sample. \**P* < 0.05; \*\**P* < 0.01. n.s. non-significant. Normality and Lognormality Tests were used to determine whether the data fit the normal distribution, two-tailed paired t-test is employed when the data follows a normal distribution, whereas the non-parametric two-tailed Wilcoxon signed rank test is utilized in cases where the data does not conform to a normal distribution.

We further identified molecularly-defined PY1 and PY2 neurons that are morphologically reconstructed by biocytin straining after patch-clamp recording (Figure 3B and Supplemental Figure 3A). We found higher transcript densities of *CDKN1A, NFKBIA* and *NEFM* in PY2 neurons (n = 7) compared with PY1 neurons (n = 7. Supplemental Figure 4). Our results showed no significant differences of morphological, intrinsic membrane characteristics or action potential firing characteristics between PY1 and PY2 neurons (Supplemental Figure 5). Through analysis of 7,688 neurons identified by ESAI-FISH from four independent brain samples, we found increased mRNA counts of *NFKBIA, CDKN1A, CCL2, CUX2, NEFM* and *NEFH* genes in molecularly-defined PY2 neurons (n = 2369) compared with PY1 neurons (n = 5319, 7 brain slices from 4 individual samples. Supplemental Figure 3H). In addition, the density of PY2 marker gene *CDKN1A* and *NFKBIA,* neurofilament marker genes *NEFM* and *NEFH,* as well as soma volume are substantially increased in molecularly-defined PY2 neurons compared with PY1 neurons (Figure 3I). Together, the molecularly-defined PY2 neurons exhibited enlarged soma size, upregulation of neurofilament genes and cellular senescence genes, suggesting that these neurons are tightly associated with pathological neurons in the brain tissue from people with drug-resistant epilepsy.

### Candidate markers and senescent state of histopathological neurons in epilepsy

The histopathological dysmorphic neurons in drug-resistant epilepsy patients with focal cortical dysplasia (FCD) exhibited enlarged cell soma size, increased expression of the non-phosphorylated neurofilament protein (SMI) and upregulation of the mTOR pathway (2, 5), reminiscent of morphological features of senescent cells(32, 33). Bulk RNA sequencing or microarray analysis reveal gene expression changes in mTOR pathway, neural development, synaptic plasticity, neuroinflammation, and neurodegeneration in the brain tissue of drug-resistant epilepsy patients or rodent epilepsy models (5, 12, 34–36). The expression of genes associated with the chemokine and inflammation pathways have been shown in cortical neurons of the epilepsy patients (11, 34, 37, 38). However, the cell type specific gene expression in the brain tissue of people with drug-resistant epilepsy remains unresolved.

To examine whether the marker genes of PY2 neurons are expressed in histopathological neurons, we performed immunohistochemistry to examine whether cell cycle inhibitor P21 (encoded by the *CDKN1A* gene) and senescent markers are expressed in histopathological dysmorphic neurons in FCD. Our results showed the expression of P21 in non-phosphorylated neurofilament protein (SMI) expressing histopathological dysmorphic neurons in the brain tissue of FCD (Figure 4A and Supplemental Figure 6, A and B). Furthermore, we found the chemokine C-C motif chemokine ligand 2 (CCL2, encoded by the *CCL2* gene) and the nuclear factor-kappa-B-inhibitor alpha (NFKBIA, encoded by the *NFKBIA* gene) are expressed in SMI positive histopathological dysmorphic neurons in the brain tissue of FCD (Figure 4A and Supplemental Figure 6, A and B). In addition, we found the expression of P21, CCL2, NFKBIA are substantially enriched in SMI expressing histopathological neurons compared with NeuN positive neurons in brain tissue in FCD (Figure 4, A and B), but not expressed in cortical neurons in control brain tissue with no history of epilepsy (Figure 4, C and D, and Supplemental Figure 6, C and D). Together, these results demonstrated that the histopathological dysmorphic neurons in drug-resistant epilepsy patients with FCD expressed PY2 marker genes that are associated with brain inflammation and cellular senescence.

**Figure 4.**
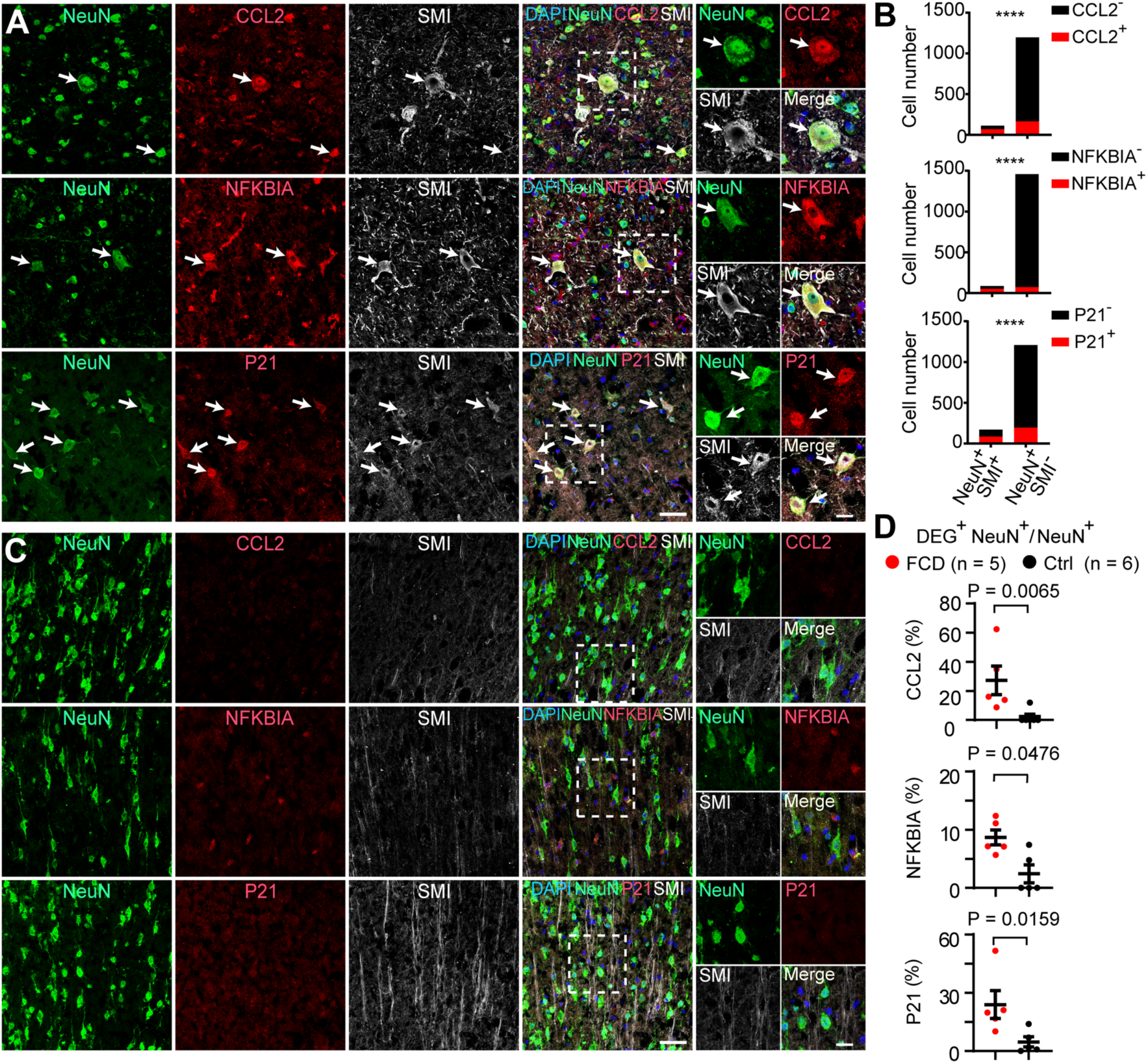
Candidate marker genes for histopathological dysmorphic neurons in drug-resistant epilepsy with focal cortical dysplasia. **(A)** Representative images and statistical quantification (**B**) showing the expression of differentially expressed genes (DEGs) in SMI-positive dysmorphic neurons compared with SMI-negative neurons in focal cortical dysplasia. Arrows indicated co-expression of PY2 marker genes with histopathological neurons. Data are presented as cell numbers quantified from 5 individual samples for each marker. Fisher’s exact-test. \*\*\*\**P* < 0.0001. **(C)** Representative images and statistical quantification (**D**) showing the expression of CCL2 (FCD: 27.2±9.9%, n = 5; Control: 2.0±2.0%, n = 6), NFKBIA (FCD: 8.7±1.3%, n = 5; Control: 2.4±1.5%, n = 5) and P21 (FCD: 23.9±7.2 %, n = 5; Control: 4.7±2.6%, n = 5) in total neurons in FCD compared with control cortical brain tissue with no history of epilepsy. Data are presented as mean ± SEM. Two-tailed Mann-Whitney test.

### Senescent neurons are the major pathological characteristics in the epileptic focus

Senescent cells are in permanent cell cycle arrest that prevents uncontrolled cell division. Recently, molecular markers for cellular senescence were identified in postmitotic cells such as neurons in nerve injury or in the neurodegenerative diseases (32, 33, 39). Cellular senescence can be induced by oxidative stress, DNA damage, or hyperactive mTOR pathway (33, 39, 40). The PY2 pyramidal neuron cluster that showed genes enrichment for inflammatory response and cellular senescence are present in brain samples from drug-resistant epilepsy patients with different pathologies (Figure 1, D and G, and Supplemental Figure 1A). Therefore, we went on to examine the senescent cell markers expression in the brain tissue of people with drug-resistant epilepsy. β-galactosidase activity is widely used to label senescent cells both *in vitro and in vivo* (23). We found senescence-associated β-galactosidase (SA-β-Gal, encoded by the *GLB1* gene) positive cells with neuronal morphology in the brain tissue of drug-resistant epilepsy with different pathologies, including FCD (14.4 ± 5.5 cells/mm^2^, n = 35), TLE (19.0 ± 8.8 cells/mm^2^, n = 17), and PT (12.1 ± 3.6 cells/mm^2^, n = 11), but not in control cortical tissue of brain samples with no history of epilepsy (n = 4. Figure 5, A-C, and Supplemental Figure 8).

**Figure 5.**
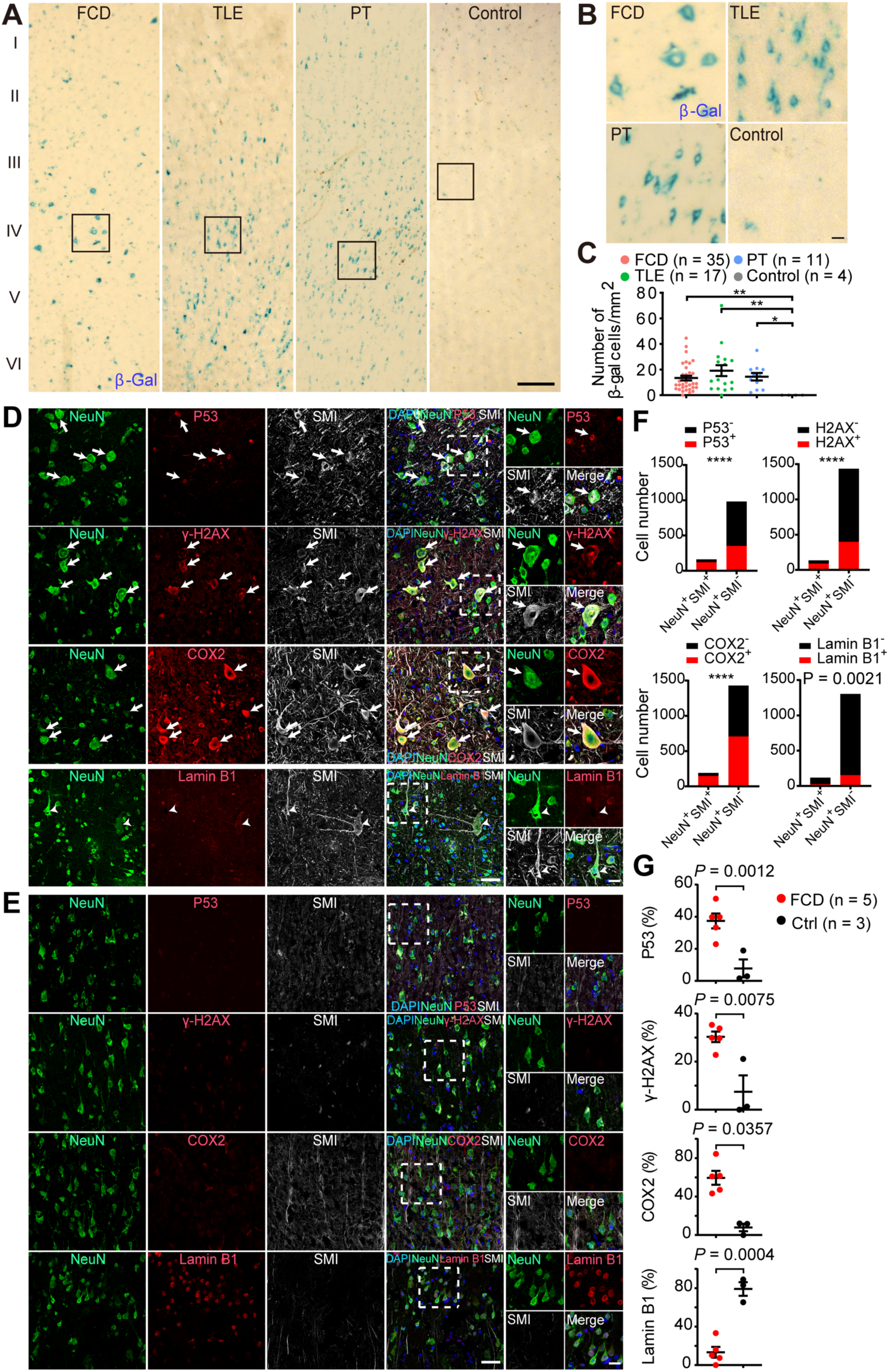
Expression of senescent cell markers in histopathological neurons in people with drug-resistant epilepsy. (**A**-**C)** Representative images (**A**) and statistical quantification (**C**) of SA-β-Gal staining showed β-Gal positive cells were present in the epileptic focus of FCD (n = 35 samples), TLE (n = 17 samples), PT (n = 11 samples), but not in control cortical tissues with no history of epilepsy (n = 4 samples). Scale bars: 200 μm (A), 20 μm in zoom-in images (B) of box area in panel (A). Data are presented as mean ± SEM. Kruskal-Wallis test with Dunn’s multiple comparisons test. \*\**P* < 0.01, \**P* < 0.05. **(D-G)** Representative images (**D**) and statistical quantification (**F**) showing the expression of marker genes in SMI-positive dysmorphic neurons compared with in SMI-negative neurons in focal cortical dysplasia or control brain samples (**E** and **G**). Arrows indicated co-expression of senescent cell markers with SMI and NeuN. Arrowheads indicated no Lamin B1 expression were detected in SMI-positive histopathological dysmorphic neurons. Data are presented as cell numbers quantified from 5 individual samples for each marker (**F**. Fisher’s exact-test. \*\*\*\**P* < 0.0001). Data are presented as mean ± SEM (**G.** Two-tailed Mann-Whitney test). Scale bar, 50 μm; 10 μm for zoom-in images.

Next, we examined the expression of senescent cell markers in the brain tissue of people with drug-resistant epilepsy compared with control brain samples. Cyclooxygenase-2 (COX2) (encoded by the *PTGS2* gene) expression can be induced by seizure activity (41, 42) and is associated with age-dependent neuronal pathology (43, 44). The *PTGS2* is upregulated in both PY1 and PY2 pyramidal neurons. We found COX2 is highly expressed in cortical neurons and enriched in SMI positive histopathological dysmorphic neurons in FCD (Figure 5, D and F, and Supplemental Figure 6, A and B), suggesting extensive oxidative stress in neurons. Cell cycle inhibitors P53 (encoded by the *TP53* gene) and DNA damage marker γ-H2AX (histone H2AX, encoded by the *H2AX* gene) are expressed in SMI positive neurons in FCD (Figure 5, D and F, and Supplemental Figure 6, A and B). In addition, the expression of P53, γ-H2AX, COX2 are substantially enriched in SMI expressing histopathological neurons compared with NeuN positive neurons in brain tissue in FCD (Figure 5, D and F), but not expressed in cortical neurons in control brain tissue with no history of epilepsy (Figure 5, E and G, and Supplemental Figure 6). Loss of Lamin B1 (encoded by the *LMNB1* gene) results in disruption of nuclear integrity and premature senescence (23, 45). We found Lamin B1 expression was downregulated in histopathological dysmorphic neurons in FCD compared with control cortical neurons (Figure 5, D-G, and Supplemental Figure 6), suggesting disrupted nuclei integrity of dysmorphic neurons in FCD.

We further demonstrated that senescent cell markers P53, P21, γ-H2AX, COX2 are highly expressed in cortical neurons in the brain tissue of drug-resistant epilepsy with different pathologies, but not in control brain tissues with no history of epilepsy (Figure 5, D-G, Figure 6, A-F, and Supplemental Figures. 6 and 7). In addition, we found substantial reduction of nuclear integrity marker Lamin B1 expression in neurons in the epileptic focus of FCD, PT, TLE brain samples in comparison with control brain tissue (Figure 5, D-G, Figure 6, E and F, and Supplemental Figures 6 and 7). These results demonstrate that cortical pyramidal neurons in the brain tissue of people with drug-resistant epilepsy exhibited hallmark features of cellular senescence, including expression of senescent markers P53, P21, COX2, γ-H2AX, SA-β-Gal and reduction of Lamin B1. The accumulation of senescent neurons in cortical brain tissue suggests a tight association between epileptic activity and neuronal senescence in people with drug-resistant epilepsy.

**Figure 6.**
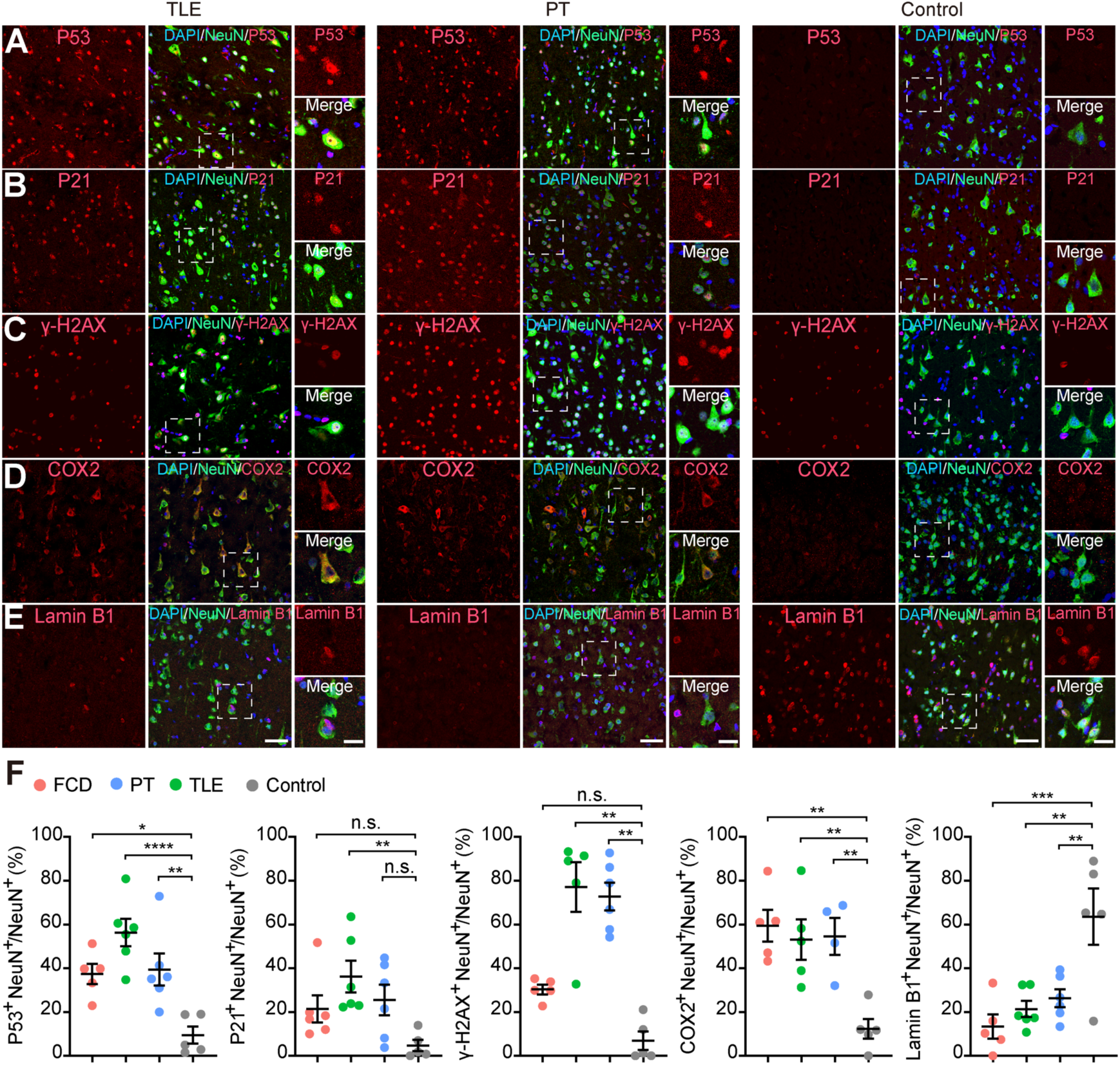
Expression of senescent markers in pathological neurons in the epileptic focus of drug-resistant epilepsy patients. (**A**-**E)** Representative images showing the co-expression of senescent cell marker genes P53 (**A**), P21 (**B**), γ-H2AX (**C**) and COX2 (**D**) with NeuN in the epileptic focus of drug-resistant epilepsy with TLE, PT and control brain samples. **(E)** Lamin B1 was downregulated in neurons in the epileptic focus of TLE, PT compared with cortical neurons in control brain samples. Scale bar, 50 μm; 10 μm for zoom-in images. **(F)** Statistical results showing the percentage of senescent marker genes expression in cortical neurons in the epileptic focus of drug-resistant epilepsy with different pathologies compared with cortical neurons in control brain samples with no history of epilepsy. P53^+^NeuN^+^/NeuN^+^ (FCD: 37.4±4.2%, n = 5; TLE: 56.3±6.2%, n = 6; PT: 39.4±7.3%, n = 6; Control: 9.5±3.6%, n = 5. *P* = 0.0003); P21^+^NeuN^+^/NeuN^+^ (FCD: 21.5±6.2%, n = 6; TLE: 36.2±7.2%, n = 6; PT: 25.5±7.0%, n = 6; Control: 4.7±2.4%, n = 5. *P* = 0.0225); γ-H2AX^+^NeuN^+^/NeuN^+^ (FCD: 30.3±2.0%, n = 5; TLE: 77.1±10.4%, n = 5; PT: 72.8±6.4%, n = 6; Control: 6.9±3.8%, n = 5. *P* < 0.0001); COX2^+^NeuN^+^/NeuN^+^ (FCD: 59.5±7.2%, n = 5; TLE: 53.1±9.2%, n = 5; PT: 54.6±8.4%, n = 4; Control: 12.3±4.5%, n = 5. *P* = 0.0013); Lamin B1^+^NeuN^+^/NeuN^+^ (FCD: 13.4±5.1%, n = 5; TLE: 21.4±3.7%, n = 6; PT: 26.4±4.1%, n = 6; Control: 63.6±11.7%, n = 5. *P* = 0.0006). Data are presented as mean ± SEM. Kruskal-Wallis test with Dunn’s multiple comparisons test. \**P* < 0.05, \*\**P* < 0.01, \*\*\**P* < 0.001, \*\*\*\**P* < 0.0001. n.s. non-significant.

**Figure 7.**
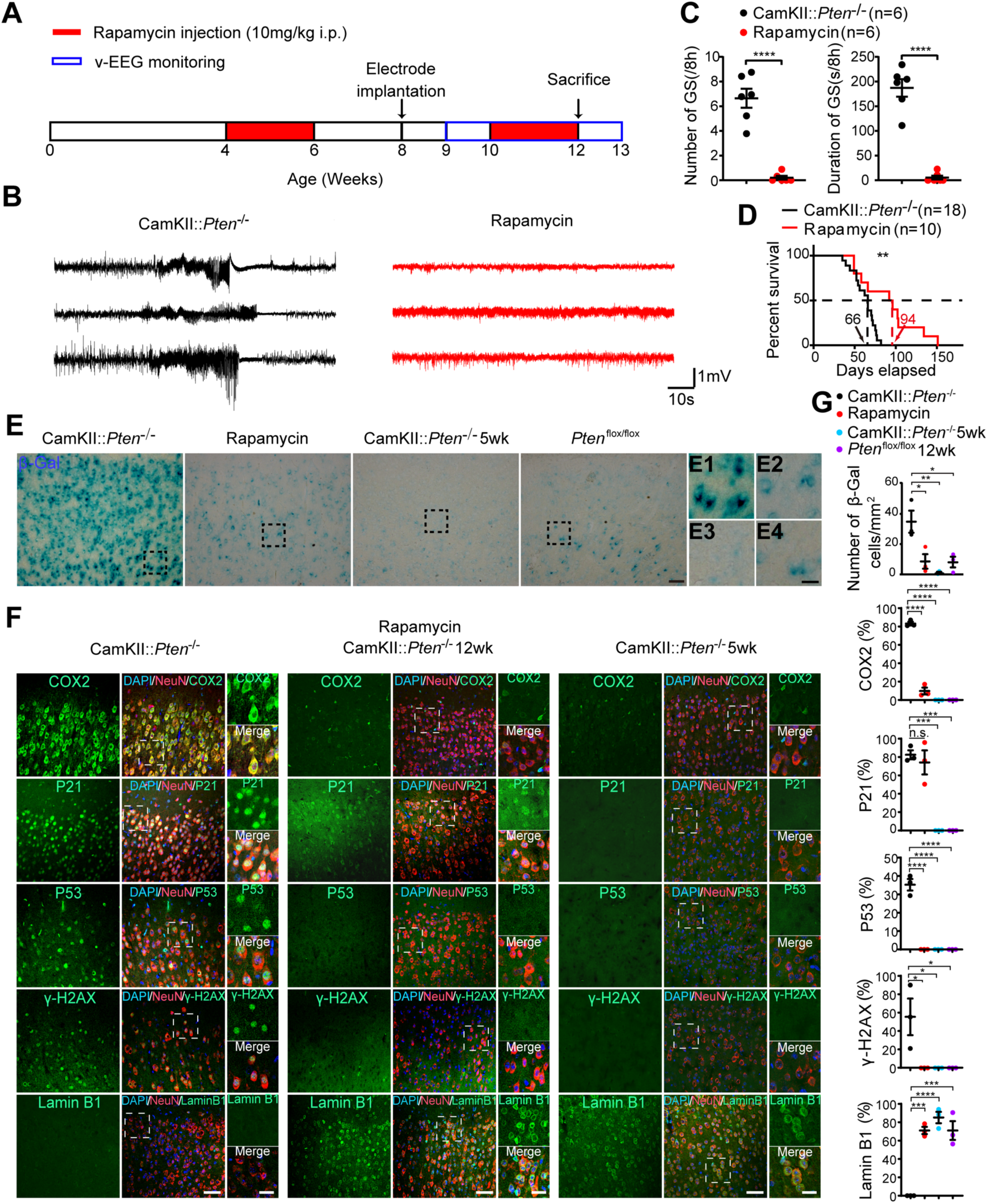
Neuronal senescent markers are induced by epileptic activities in mouse model of drug-resistant epilepsy. **(A)** Schematic diagram showing the experimental design of video EEG recording and rapamycin injection (i.p.) of CamKII::*Pten*^-/-^ mice. **(B)** Representative traces showing generalized seizures (G.S.) from EEG recordings of vehicle (black traces, n = 3 mice) and rapamycin-treated (red traces, n = 3 mice) CamKII::*Pten*^-/-^ mice. **(C)** Average number and duration of generalized seizures (GS) were substantially reduced in rapamycin-treated compared with vehicle-treated CamKII::*Pten*^-/-^ mice (n = 6 mice per group). \*\*\*\**P* < 0.0001. Two-tailed *t*-test. **(D)** Rapamycin treatment Supplemental the lifespan of CamKII::*Pten*^-/-^ mice. Black trace: vehicle-treated group (n = 18 mice); red trace: rapamycin-treated group (n = 10 mice). *P* = 0.0047, Log-rank (Mantel-Cox) test. **(E** and **F)** Representative images showing the expression of senescent markers (SA-β-Gal, COX2, P21, P53, γ-H2AX) and reduction of nuclear integrity marker Lamin B1 in CamKII::*Pten*^-/-^ mice with status epilepticus, but not in CamKII::*Pten*^-/-^ mice that had not developed generalized seizures at 5 postnatal week (5wk), or in control *Pten*^flox/flox^ mice at 12wk. Rapamycin treatment repressed senescent markers expression in CamKII::*Pten*^-/-^ mice. Expression of Lamin B1 can be restored after rapamycin treatment of CamKII::*Pten*^-/-^ mice. Arrows indicated co-expression of senescent markers with NeuN. Scale bar, 50 μm; 10 μm for zoom-in images. **(G)** Statistical results showing the proportion of senescent markers and NeuN co-expressing neurons in total cortical neurons. n = 3 mice per group. Data are presented as mean ± SEM. Kruskal-Wallis test with Dunn’s multiple comparisons test. \**P* < 0.05, \*\**P* < 0.01, \*\*\**P* < 0.001, \*\*\*\**P* < 0.0001. n.s. non-significant.

### Chronic epileptic seizures induce neuronal senescence in mouse models of drug-resistant epilepsy

Next we examined whether neuronal senescent markers expression can be induced by epileptic activities in mouse models of drug-resistant epilepsy. *PTEN* deficient neurons exhibit hyperactivation of mTOR pathway and pathological features resembling histopathological neurons in drug-resistant epilepsy that cause epileptic seizures (46, 47). The transgenic CamKII-cre mice with selective deletion of *Pten* (CamKII::*Pten*^-/-^) developed generalized tonic-clonic seizures at 6 postnatal weeks and died by 9 postnatal weeks on average (Figure 7, A-C). We found upregulation of senescent markers SA-β-Gal, P53, P21, γ-H2AX, COX2 and downregulation of Lamin B1 in cortical pyramidal neurons in CamKII::*Pten*^-/-^ mice with status epilepticus, but not in CamKII::*Pten*^-/-^ mice that had not developed seizures at 5 postnatal weeks or control *Pten*^flox/flox^ mice at 12 postnatal weeks (Figure 7, D-F). We also established acute and chronic epilepsy model by focal intra-hippocampal injection of kainic acid. We found the expression of senescent markers SA-β-Gal, P21, γ-H2AX and COX2 as well as downregulation of Lamin B1 in cortical neurons in mice that developed chronic focal seizures and generalized tonic-clonic seizures two months after kainic acid injection (Supplemental Figure 10), but not in mice 3 days post acute epileptic seizures (Supplemental Figure 9). These results suggest that chronic epileptic seizures can induce neuronal senescence in mouse models of drug-resistant epilepsy.

We further examined the expression of senescent markers in cortical neurons after suppressing epileptic seizures. Chronic treatment of rapamycin, an mTOR pathway inhibitor, suppressed generalized tonic-clonic seizure frequency and Supplemental the lifespan of CamKII::*Pten*^-/-^ mice (Figure 7, A-C). In addition, we found substantial downregulation of senescent neuronal markers SA-β-Gal, P53, COX2 and γ-H2AX, as well as restoration of Lamin B1 expression in cortical pyramidal neurons in CamKII::*Pten*^-/-^ mice at 12 postnatal weeks following rapamycin treatment (Figure 7, D-F). The P21 expression was relatively weaker but the percentage of cortical neurons expressing P21 was not changed in CamKII::*Pten*^-/-^ mice after rapamycin treatment compared with P21 expression in CamKII::*Pten*^-/-^ mice with status epilepticus (Figure 7, E and F). Together, these findings demonstrated the repression of neuronal senescent markers by blocking epileptic seizures in rodent models of drug-resistant epilepsy.

To summarize, our results revealed that cortical pyramidal neurons exhibited hallmarks of cellular senescence in the epileptic focus of people with drug-resistant epilepsy. The expression of senescent cell markers is induced by chronic epileptic seizures and tied to the pathological feature of histopathological neurons in drug-resistant epilepsy. These molecular markers for neuronal senescence are important for the histopathological diagnosis and provide a new perspective in uncovering the mechanism underlying the pathogenesis of drug-resistant epilepsy.

## Discussion

### Patch-seq analysis revealed molecular features of histopathological neurons in human drug-resistant epilepsy

In the present study, we used patch-seq analyses that integrate whole-cell electrophysiology with single-cell RNA sequencing to reveal the molecular features of pathological neurons in the brain tissue from people with drug-resistant epilepsy. The 197 patch-seq neurons analyzed in our study were collected from 36 individual donor samples and PY2 pyramidal neurons composed of 13.7% of total patch-seq neurons (27 PY2 out of 197 total patch-seq neurons) and 18.8% of total pyramidal neurons (27 PY2 out of 144 PY patch-seq neurons. Figure 1C). These results are consistent with previous findings that histopathological dysmorphic neurons are cortical pyramidal neurons (28, 48) and suggest that patch-seq approach enables us to enrich pathological neurons for single-cell analysis in brain tissue from people with drug-resistant epilepsy. We further performed variation partition analysis to assess genes associated with individual variation (49) and found that the majority of genes expression showed highest variability associated with subtype, while exhibited low variability associated with donor, gender, brain region, age, layer and pathology (Supplemental Figure 2, A and B). In addition, there is substantial overlap between subtype marker genes and high variance genes associated with subtype (> 20% variance explained), but not with high variance genes associated with donor (> 20% variance explained). The transcriptomic cell subtypes showed a lack of correspondence with pathology, donor, brain area, cortical layer distribution, age and gender as shown in the UMAP plots (Supplemental Figure 2, C and D). These results suggest that the identification of subtype-specific markers was not affected by donor variance.

We compared the expression of PY2 transcriptomic profiles in brain tissue from epilepsy patients examined with control brain tissue with no history of epilepsy from public available single-nucleus RNA sequencing dataset (49, 50). Our result shows enrichment of PY2-like cells in cortical pyramidal neurons in brain samples from epilepsy patients compared with non-epilepsy controls (Supplemental Figure 1, C-H). In addition, the mTOR pathway genes RPS6, TPI1, ENO1 and PGK1 are enriched in PY2-like cortical neurons of brain samples from epilepsy patients compared with non-epilepsy controls. Although previous studies reported pro-inflammatory signaling in brain tissue from epilepsy patients (50), our results revealed that PY2 pyramidal neurons exhibited transcriptomic signatures including upregulation of genes associated with mTOR pathway, cellular senescence and the inflammatory response pathway, mimicking the molecular features of histopathological neurons in drug-resistant epilepsy.

### Integrated analyses reveal the molecular and morphological features of pathological neurons in the brain tissue of people with drug-resistant epilepsy

Despite substantial changes in gene expression profiles, the PY2 pyramidal neurons exhibited similar intrinsic membrane properties and action potential firing characteristics in comparison with the PY1 pyramidal neurons. The PY3 pyramidal neurons demonstrated smaller membrane capacitance (Cm) and higher instant AP firing frequency compared with PY1 and PY2 pyramidal neurons (Figure 2, A and C) (51, 52). However, PY3 are a small group of pyramidal neurons recorded from the temporal lobe (Figure 1G), possibly representing characteristics of regional subpopulation of pyramidal neurons. There are 39.2% of patch-seq neurons (127 out of total 324 neurons) that did not pass the quality control for subsequent single-cell RNA sequencing analysis. The success rate of retrieving the morphology of recorded neurons and extracting cell soma for single-cell RNA sequencing was also very low. Retrieving the full dendritic morphology of patch-seq neurons requires Supplemental loading time of biocytin, but most RNA sequencing data from morphologically reconstructed dysmorphic neurons did not pass the quality control, possibly due to that prolonged whole-cell recording time resulted in mRNAs degradation and low yield of cDNA (53).

We therefore developed the state-of-the-art approach that integrated patch-clamp recording with the EASI-FISH staining to simultaneously examine multiple marker genes expression in single neurons in thick brain slices after electrophysiological recording. By analyzing 7,688 cortical pyramidal neurons from four individual brain samples of drug-resistant epilepsy, our results reveal molecularly-defined PY1 and PY2 neurons that exhibit distinct gene expression profiles. The PY2 neurons also exhibited increased expression of neurofilament genes and larger soma volume that are characteristic of pathological neurons in drug-resistant epilepsy. This EASI-FISH approach enables us to characterize the soma volume of molecularly-defined cell subtypes at scale and will thus be more efficient in identifying and characterizing the molecular features of senescent cells in tissue.

We found no significant differences in electrophysiological or morphological parameters of molecularly-defined PY1 and PY2 neurons that were also morphologically reconstructed by biocytin staining. The power of statistical analysis could be limited due to the relatively small sample size when analyzing the morphological and electrophysiological characteristics of biocytin-loaded PY1 (n = 7) and PY2 neurons (n = 7. Figure 3 and Supplemental Figures 3-5). This limitation stemmed from the scarcity of suitable human brain samples for electrophysiological recordings, the challenges of reconstructing the morphology of recorded neurons, and the time-consuming, technically demanding process of multiplex EASI-FISH analysis on neurons in the thick brain slices after patch-clamp recording.

These results may suggest that neurons within the same transcriptomic subtypes exhibited variations of morphological and electrophysiological characteristics, consistent with published reports on patch-seq analysis of both human and mouse cortical neurons (20, 54). One potential explanation could be that changes in senescent gene expression preceded the changes in morphological or electrophysiological characteristics in pathological neurons. Previous studies from both our group and others showed enhanced GABAergic synaptic activity and upregulation of the Na^+^-K^+^-Cl^-^ co-transporter (NKCC1) in pathological neurons, suggesting the GABAergic activity in pathological neurons could be depolarizing (28, 55, 56). The breakdown of cortical inhibitory restraint and propagation of excitatory glutamatergic activity, and subsequently lead to the initiation and spread of seizure activity (57, 58). Therefore, the imbalance of synaptic excitation and inhibition of pathological neurons could contribute to seizure genesis.

### The expression of senescent neuronal markers is tightly associated with epilepsy

Cellular senescence plays important roles in embryonic development, aging, and is also implicated in various diseases (23, 24). Recent studies have demonstrated the expression of senescent cell markers in retinal ganglion neurons, dopamine neurons and Purkinje neurons upon nerve injury or in the aging brain (39, 59, 60). Our results revealed the expression of senescent cell markers P21, P53, COX2, γ-H2AX, SA-β-Gal and reduction of the nuclear integrity marker Lamin B1 in SMI-expressing histopathological neurons in drug-resistant epilepsy patients with different pathologies, but not in control brain tissue with no history of epilepsy. We further established mouse models of drug-resistant epilepsy and found that neuronal senescence can be induced in mice that developed chronic epileptic seizures (Figure 7 and Supplemental Figure 10), but not in cortical neurons in mice 3 days post intra-hippocampal injection of kainic acid (Supplemental Figure 9). These results demonstrated that chronic epileptic seizures can induce neuronal senescence in drug-resistant epilepsy.

Generalized tonic-clonic seizure activity might cause vasocontractions and reduced blood and oxygen supply, that might subsequently cause prolonged stroke-like hyperoxia in the epileptic focus (42, 61). Upregulation of COX2 during hypoxia in the postictal phase of epilepsy is associated with postictal memory and behavioral deficits (42, 61). Severe hypoxia in the seizure onset zone can induce upregulation of COX2 that is closely correlated with cognitive, sensory and motor impairments in patients with chronic epilepsy (42, 61). Accumulated DNA damage marker γ-H2AX in postmitotic neurons might induce the expression of cell cycle dependent genes and genome instability of neurons (23, 62). Upregulation of cell cycle inhibitor P21 plays an important role in neuronal senescence (32, 33, 39). The P53 tumor-suppressor gene can be induced by excitotoxic stimulation and is responsible for neuron viability (63–65). Programmed cellular senescence is dependent on P21, but independent of P53 (66, 67). These results suggested that expression of the senescent marker genes may contribute to the pathogenesis of histopathological neurons in drug-resistant epilepsy (40). The senescent cells might induce morphological alterations and senescent markers expression of neighboring pyramidal neurons through the secretory senescence-associated secretory phenotype (SASP) factors and are therefore important targets for treating drug-resistant epilepsy or neurodegenerative diseases (32, 40).

Taken together, by integrating a series of state-of-the-art approaches to analyze cortical pyramidal neurons, our study revealed molecular characteristics and senescent state of pathological neurons in the brain tissue from people with drug-resistant epilepsy. These results shed new light on the pathophysiological mechanism of drug-resistant epilepsy and the development of potential therapeutic approaches targeting senescent neurons.

## Methods

Detailed information on materials and methods is provided in Supplemental Material.

### Sex as a biological variable

The human brain samples used in this work were collected from male and female individuals (Supplemental table 1). Moreover, our study examined male and female animals, and similar findings are reported for both sexes.

### Tissue collection

Human brain tissue was obtained from the surgical resections of patients with medically intractable epilepsy at the Second Affiliated Hospital of Zhejiang University and National Health and Disease Human Brain Tissue Resource Center. All human brain tissue underwent pathological examination and 78 specimens in total were collected for sequencing and immunostaining, the detailed sample information was listed in Supplementary table 1.

### Animals

CamKII-cre mice (The Jackson Laboratory, 005359) were crossed with *Pten*^flox/flox^ mice (The Jackson Laboratory, 006440) to generate CamKII-cre::*Pten*^-/-^ mice.

### Patch-seq

Strict steps were taken to avoid potential environmental RNases contamination. Surface area of electrophysiological rig was thoroughly cleaned with 70% ethanol and RNaseZap (Invitrogen, AM9780) before the experiment. Borosilicate glass electrodes (Sutter, BF150-86-10) were previously wrapped with thin foil and sterilized at 180°C in an oven for two hours. Recording electrodes (3-6 MΩ) were made on P97 pipette puller (Sutter Instrument), and filled with less than 3 μl internal solution. Immediately after electrophysiological recording, a slight negative pressure was applied to aspirate the cell cytosol and the nucleus. After the nucleus was attracted to the pipette tip, the patch electrode was retracted slowly until a globule was seen at the tip of the patch electrodes, indicating successful harvest of the cell. The tip of patch electrode was broken in a PCR tube containing 4 μl cell lysis buffer, which contains dNTPs, Oligo (dT) primer, RRI and ERCC, to ensure full capture of the cell contents in internal solution. The PCR tube was then immediately stored at −80°C.

The cDNA amplification for individually harvested cells was performed following the Smart-seq2 protocol. Briefly, mRNA in the harvested cell was denatured at 72°C for 3 min in lysis buffer. Denatured RNA was reversely transcribed into cDNA. cDNA was then pre-amplified with ISPCR primers and KAPA HotStart ReadyMix (KAPA, kk2602) for 21 PCR cycles. After amplification, only cells with a cDNA yield greater than 1 ng and appropriate size distribution (main peak at 1000-5000 bp) was retained for subsequent library construction. We used ERCC RNA spike-in control mixes during cDNA library preparation and assess the sample quality of full-length cDNA profiles of individual patch-seq neurons by an Bioanalyzer 2100 (Agilent) or Qseq100 (BiOptic). Final patch-seq neuron cDNA libraries were prepared from 1 ng of cDNA using the True Prep TM DNA Library Prep Kit V2 (Vazyme, TD-503-01). Then the cDNA libraries were sequenced in the paired-end mode on a HiSeq 1500 platform to approximately 2 million reads per cell.

### EASI-FISH

Following whole-cell patch-clamp recordings, human brain slices were post-fixed in 4% paraformaldehyde (PFA) in 1x PBS for 10-12 hours. Subsequently, slices were washed twice with 1x PBS and stored in 70% ethanol for up to 3 weeks. Before standard EASI-FISH processing, the slices were rehydrated in 1x PBS and incubated overnight with 1:1000 Alexa Fluor 555-conjugated streptavidin diluted in 1x PBS with 0.1% Tween-20 (PBST). Following incubation, the slices were rinsed twice with 1x PBS for 20 minutes each. The slices were then mounted onto glass-bottom dishes (Cellvis) and imaged using an FV3000 confocal microscopy (Olympus) equipped with a 20x/1.0 objective. Regions of interest (ROIs) were identified and imaged. After image acquisition, the brain tissue was dissected under an MVX10 stereo microscopy (Olympus). ROIs were carefully dissected into smaller pieces (1-4 mm^2^) containing the recorded cells.

The small dissected tissues underwent RNA anchoring, gelation, Proteinase K digestion and DNase I digestion. Subsequently, 3 rounds of HCR RNA-FISH were performed to detect the expressions of 6 candidate marker genes (*NFKBIA, CDKN1A, NEFM, CCL2, CUX2, NEFH*), with 2 genes in each round, because the 546-channel was occupied by streptavidin. Prior to imaging, the samples were stained with 5 µg/ml 4’,6-diamidino-2-phenylindole (DAPI) for 2 x 30 minutes. All samples were then imaged in PBS with 2x expansion using a Zeiss Lightsheet 7 microscope. Imaging was conducted using a 20x water-immersion objective (20x/1.0 W Plan-Apochromat Corr DIC M27 75 mm, RI = 1.33) with 1x zoom. Images were collected at a pixel resolution of 0.23 x 0.23 µm (post-expansion) and a z-step size of 0.4 µm with single camera detection across four tracks: the 405 nm, 488 nm, 546 nm and 669 nm channels. After image acquisition, the probes and HCR hairpins were removed using DNase I.

### Statistical analysis

We used different statistical analysis approaches to handle potential sources of individual variability due to variation of sample size in different experiments. The differential gene expression of single-cell RNA sequencing of 197 patch-seq neurons were collected from 36 individual human brain samples. Statistical analysis of differential gene expression was performed using the FindAllMarkers function of Seurat R package (version 4.0.3) with the default Wilcox Rank Sum test. Genes with adjust p-value < 0.05 and fold change > 1.5 were defined as subtype markers, Wilcox test was used to compare the expression levels of subtype markers between different cell subtypes. Fisher’s exact test was employed for the comparison of categorical variables, two-tailed *t*-test was used to compare continuous variables between two groups, Kruskal−Wallis test was utilized to estimate the variance among multiple groups. The p-values in multiple tests were adjusted to false discovery rate (FDR) using the Benjamini-Hochberg method. Data visualization was performed using ggplot2 and ComplexHeatmap R packages.

The marker genes expression in brain tissue sections was validated by immunohistochemistry from 5 to 7 individual human brain samples of each disease pathology. Fisher’s exact test was used to compare marker genes expression in SMI-positive dysmorphic neurons versus SMI-negative neurons in FCD. Normality and Lognormality tests were used to determine whether the data fit the normal distribution, two-tailed *t*-test is employed when the data follows a normal distribution, whereas the Non-parametric two-tailed Mann-Whitney test is utilized in cases when the data does not conform to a normal distribution. To determine significant differences between three or more groups, we used non-parametric Kruskal-Wallis test with Dunn’s multiple comparisons test. The statistical analyses were performed using Prism 7 software (Graphpad). \**P* < 0.05; \*\**P* < 0.01; \*\*\**P* < 0.001; \*\*\*\**P* < 0.0001. Data are presented as mean ± standard error of the mean (SEM).

### Study approval

Human surgical specimens were collected only with patient consent and research protocols were approved by the Committee on Human Research at Zhejiang University (I2021001618). Mice were housed and treated according to the guidelines of the Zhejiang University (ZJU) Laboratory Animal Care and Use Committee and were approved by the Animal Advisory Committee at ZJU (ZJU20210231).

### Data and Code availability

Processed cell-by-gene data and all R codes used for single cell RNA-seq analysis of patch-seq neurons were uploaded to the GitHub repository (https://github.com/FlameHuang/ZJU_DRE_Patch_seq_analysis). The sequencing data of patch-seq dataset has been deposited in BIG GSA PRJCA015942 under accession no. HRA006046 at https://www.cncb.ac.cn/. The customized Matlab (MathWorks) scripts are available at https://github.com/Saintgene-Xu-lab/Colocalization-analysis. Source data for this study are available in the supplemental Supporting Data Value file. Further information and requests for resources and reagents should be directed to and will be fulfilled by the corresponding authors.

## Author Contributions

JC conceived and designed the project, QG, XD, YZ, YS, JP, HZ performed patch-seq experiment and electrophysiological data analysis. FH, JG, YZ, LS analyzed single-cell RNA sequencing data from patch-seq neurons. XD, CC, SX performed EASI-FISH experiment and data analysis. JY, CC, MW, MZ, JS, YH, SX performed immunohistochemistry and data analysis. JY, MZ performed EEG recording, pharmacology and immunohistochemistry in mouse models of drug-resistant epilepsy. SD, YW, XL, ZC, YS, JZ, JZ provided reagents. QG, JY, FH, XD, SD, CC, SX, LS, JC analyzed data. JC supervised the project and prepared the manuscript with inputs from all authors.

## Acknowledgement

We thank Drs. Aimin Bao, Keqing Zhu, and Juanli Wu at the National Health and Disease Human Brain Tissue Resource Center at Zhejiang University and staff at the Second Affiliated Hospital of Zhejiang University for providing brain samples. We thank Zhaoxiaonan Lin and staff from the Core Facilities of Zhejiang University School of Medicine for their technical support. We thank Dr.Jeremy Miller (Allen Brain Institute, Seattle, USA) for suggestions on the analysis of public available snRNA-seq data of the human brain. We thank Dr. Hao Wang (University of Science and Technology of China, Hefei) for help and advice in multiplex FISH. We thank Mingpo Yang for help with the MatLab program for measurement of electrophysiological parameters. We thank Dr. Jinghai Chen (Zhejiang University, Hangzhou, China) and Dr. Woo-ping Ge (Chinese Institute for Brain Research, Beijing, China) for kindly providing transgenic mice and reagents. We thank Drs. Hailan Hu, Jianhong Luo, Wei Liu and Dante Neculai (Zhejiang University, Hangzhou, China), and Dr. Wenbiao Gan (Shenzhen Bay laboratory, Shenzhen, China) for critical reading and suggestions on the manuscript. This work was supported by grants from the Ministry of Science and Technology (2019YFA0110103, 2019YFA0110101, 2022YFA1103702, 2022YFA1103200), the National Natural Science Foundation of China (81870898, 32321002, 32270839, 32371072 and 32321003), the STI2030-Major Projects (2021ZD0204401), the Fundamental Research Funds for the Central Universities (2021FZZX001-37, 2019FZA7009 and 226-2024-00055), the Zhejiang Provincial Natural Science Foundation (LR18H090002, LZ23C070003), and the Non-profit Central Research Institute Fund of Chinese Academy of Medical Sciences (2023-PT310-01).

## Competing Interests

All authors claim that there are no competing interests.

## Notes

### Competing Interest Statement

The authors have declared no competing interest.

