## Supplemental Material for "Multimodal single-cell analyses reveal molecular markers of neuronal senescence in human drug-resistant epilepsy"

This file includes:

Supplemental Methods

Supplemental Figures 1 to 10

Supplemental Table 1

Supplemental Excel table titles and legends

Supplemental References

### Supplemental Methods

#### Tissue collection and processing

Human brain tissue was obtained from the surgical resections of patients with medically intractable epilepsy at the Second Affiliated Hospital of Zhejiang University and National Health and Disease Human Brain Tissue Resource Center. Surgical specimens were collected only with patient consent and research protocols were approved by the Committee on Human Research at Zhejiang University (I2021001618). All human brain tissue underwent pathological examination and 78 specimens in total were collected for sequencing and immunostaining, the detailed sample information was listed in supplementary table 1.

Tissue collection and acute brain slices were prepared as previously described (1, 2). Surgical specimens were collected shortly after surgery in modified ice-cold oxygenated (5% O2 and 95% CO2) artificial cerebral fluid (aCSF) and immediately transferred to laboratory. The aCSF solution (pH 7.3-7.4, osmolarity 300-310 Osm) contained (in mM): N-methyl-D-glucamine (NMDG) 93, KCl 2.5, $NaH_2PO_4{\cdot}2H_2O$ 1.2, $NaHCO_3$ 30, HEPES 20, Glucose 25, Na-ascorbate 5, Na-pyruvate 3, Thiourea 2, MgSO4 10, $CaCl_2{\cdot}2H_2O$ 0.5. Brain slice sections (300 μm) were prepared on a vibratome (Leica, VT1200S) in ice-cold oxygenated NMDG aCSF solution. Then brain slice sections were transferred to a chamber containing oxygenating NMDG aCSF solution placed in 32°C water bath for 30 min incubation. Immediately after the incubation, sodium spike-in solution (2M NaCl dissolved in fresh NMDG solution) was added sequentially to the incubation chamber: 250 μl at 0 min, 250 μl at 5 min, 500 μl at 10 min, 1 ml at 15 min, 2 ml at 20 min. Subsequently, brain slice sections were transferred and incubated in oxygenated HEPES aCSF solution at room temperature. HEPES aCSF solution (pH 7.3-7.4, osmolarity 300-310 Osm) containing (in mM): NaCl

92, KCl 2.5, $NaH_2PO_4 \cdot 2H_2O$ 1.2, $NaHCO_3$ 30, HEPES 20, Glucose 25, Na-ascorbate 5, Na-pyruvate 3, Thiourea 2, $MgSO_4$ 2, $CaCl_2 \cdot 2H_2O$ 2. After more than one hour incubation, the brain slice was transferred to a recording chamber continuously perfused with oxygenated recording aCSF solution at 32°C (pH 7.3-7.4, osmolarity 300–310 mOsm) containing (in mM): NaCl 125, KCl 2.5, $NaH_2PO_4 \cdot 2H_2O$ 1.3, $MgCl_2$ 1.3, $CaCl_2$ 2, $NaHCO_3$ 25, glucose 10.

**Patch-seq electrophysiological recordings and single-cell RNA sequencing**

Patch-seq was performed as previously described (3). Strict steps were taken to avoid potential environmental RNases contamination. Surface area of electrophysiological rig was thoroughly cleaned with 70% ethanol and RNaseZap (Invitrogen, AM9780) before the experiment. Borosilicate glass electrodes (Sutter, BF150-86-10) were previously wrapped with thin foil and sterilized at 180°C in an oven for two hours. Recording electrodes (3-6 MΩ) were made on P97 pipette puller (Sutter Instrument), and filled with less than 3 μl internal solution containing (in mM): K-D-gluconate 110; KCl 4, HEPES 10, Mg-Adenosine 5'-triphosphate 4, Na-guanosine 5'-triphosphate 0.3, $Na_2$-phosphocreatine 10, EGTA 0.2, glycogen 20 μg/ml, biocytin 0.5%, recombinant RNase inhibitor (RRI), 0.5 U/μl, pH 7.3-7.4, osmolarity 320–330 mOsm.

Neurons in brain slice were visualized under differential interference contrast microscopy (Olympus, BX51), the histopathological neurons were reported to be cortical pyramidal neurons in previous studies from other groups and our group (1, 4), we therefore aim to target pyramidal neurons for patch-seq analysis and also patched interneurons for comparative analysis of the transcriptomic, morphological and electrophysiological features. After giga-seal was formed between electrode tip and cell soma, short negative pressure combined with Zap were applied to induce cell membrane rupture and established whole-cell configuration. Current pulses (500 ms or 1 s in duration) were injected into the cell to elicit steady-state action potential firing under current clamp mode. Immediately after electrophysiological recording, a slight negative pressure was applied to aspirate the cell cytosol and the nucleus. After the nucleus was attracted to the pipette tip, the patch electrode was retracted slowly until a globule was seen at the tip of the patch electrodes, indicating successful harvest of the cell. The tip of patch electrode was broken in a PCR tube containing 4 μl cell lysis buffer, which contains dNTPs, Oligo (dT) primer, RRI and ERCC, to ensure full capture of the cell contents in internal solution. The PCR tube was then immediately stored at -80°C.

**Statistical analyses of electrophysiological parameters**

We used the following quality controls to filter recorded neurons for electrophysiological parameters measurement: the recorded cells that exhibited giga-seal during initial whole-cell configuration, access resistance (Ra) less than 30 MΩ after membrane rapture, and leak current smaller than 150 pA during whole-cell recordings were selected for subsequent analyses. We extracted 13 electrophysiological parameters that represent the intrinsic membrane properties of neurons and features of action potential firing pattern by Clampfit 10.7 (Molecular Device) and custom-written program in Matlab (MathWorks, R2017b). Specifically, resting membrane potential (RP) was measured immediately after membrane rupture under current clamp mode (I = 0). Rheobase denoted minimum current injection required to elicit action potential firing. Input resistance (Rin) was determined by calculating the slope of linear regression established between voltage (mV) and inject current (pA). Time constant (Tau) was determined by the time interval between stimulation initiation and 63% of maximum voltage deflection. Membrane capacitance was determined by the quotient dividing Tau by Rin. Threshold was determined by the voltage when the derivative of action potential equals to 10. Peak was determined by the exact voltage value of an action potential. Trough was determined by the minimum voltage value of an action potential. Amplitude was determined by the voltage deflection between peak and trough. Afterhyperpolarization (Ahp) was determined by the voltage deflection between threshold and trough. Halfwidth was determined by the duration of action potential when voltage equals half of the amplitude. Latency was determined by the time interval between the onset of stimulation and action potential initiation during rheobase stimulation. Upstroke was determined by the maximal derivative in the rising phase of action potential, while downstroke was determined by the minimum derivative in the descending phase of action potential. Upstroke downstroke ratio (UDR) was as the absolute quotient between upstroke and downstroke. Stepped depolarization current was injected into the neuron till the neuron elicited maximal firing rate of action potentials (the value of injected current was recorded as suprathreshold current for individual neurons). Instaneous frequency (Ins.Freq) was determined by the reciprocal value of first interspike interval (ISI) during maximum firing rate of action potentials. Frequency was determined by the maximum firing rate of action potentials. Adaptation index (AI) was calculated by the following formula: $adaptation\ index = \frac{1}{N-1}\sum_{i=1}^{N-1}\frac{ISI_{i+1}-ISI_{i-1}}{ISI_{i+1}+ISI_{i-1}}$.

**Patch-seq library preparation and data analyses**

The cDNA amplification for individually harvested cells was performed following the Smart-seq2 protocol (5). Briefly, mRNA in the harvested cell was denatured at 72°C for 3 min in lysis buffer. Denatured RNA was reversely transcribed into cDNA. cDNA was then pre-amplified with ISPCR primers and KAPA HotStart ReadyMix (KAPA, kk2602) for 21 PCR cycles. After amplification, only cells with a cDNA yield greater than 1 ng and appropriate size distribution (main peak at 1000-5000 bp) was retained for subsequent library construction(3). We used ERCC RNA spike-in control mixes during cDNA library preparation and assess the sample quality of full-length cDNA profiles of individual patch-seq neurons by an Bioanalyzer 2100 (Agilent) or Qseq100 (BiOptic). Final patch-seq neuron cDNA libraries were prepared from 1 ng of cDNA using the True Prep TM DNA Library Prep Kit V2 (Vazyme, TD-503-01). Then the cDNA libraries were sequenced in the paired-end mode on a HiSeq 1500 platform to approximately 2 million reads per cell.

The raw sequencing reads were aligned to GRCh38 (hg38) using HISAT2 (v.2.2.1). Samtools (v1.11) was used to calculate alignment rate and remove PCR duplicates and only uniquely aligned reads were used for gene quantification. Exonic and intronic reads were used to calculate gene abundance using featureCounts (Subread v2.0.1) with an annotation retrieved from Gencode (https://www.gencodegenes.org/human/release_40.html). We applied stringent quality control procedures for analyzing the single-cell RNAseq data of patch-seq neurons. We filtered cells based on the following criteria: cells with an uniquely alignment rate < 60%, detected mRNA molecules < 70,000, the number of expressed genes < 4,000 and the percentage of mitochondrial genes > 20% were discarded, leaving 197 cells (out of total 324 cells) for further analysis. The cell-by-gene count matrix was subsequently analyzed using Seurat (v4). Gene expression levels were calculated as counts per million reads (CPM). For cell clustering, we used the 2000 most variable genes to perform principle component analysis (PCA), and selected the top 8 PCs for Uniform Manifold Approximation and Projection (UMAP) analysis. Fisher’s exact test was employed for the comparison of categorical variables. Four cell subtypes were identified based on the Louvain-clustering algorithm with the resolution of 0.6. Differentially expressed genes between subtypes were identified using the Wilcoxon Rank Sum test of the “FindMarkers” function. The p-values in multiple tests were adjusted to false discovery rate (FDR) using the Benjamini-Hochberg method. Kruskal−Wallis test was utilized to estimate the variance among multiple groups. All statistical analyses were performed using R software (version 4.0.3).

**Gene enrichment analysis**

Gene ontology (GO) analysis of subtype marker genes was performed using clusterProfiler R package based on the biological process (BP) database. Gene set enrichment analysis (GSEA) was carried out to identify the upregulated pathways in the PY2 subtype. “HALLMARK_MTORC1_SIGNALING” and “HALLMARK_INFLAMMATORY_RESPONSE” pathway genes were obtained from the human Hallmark database. Cellular senescence genes were extracted from the “FRIDMAN_SENESCENCE_UP” term of molecular signatures database (MSigDB).

**Variation partition analysis**

To account for the gene variance across different phenotype variables and clinical traits, we performed the variation partition analysis using the variancePartition R package. A linear mixed-effect model was defined as: gene ~ age + (1|donor) + (1|sex) + (1|subtype) + (1|brain region) + (1| layer) + (1|pathology). Based on the fitted variation partitioning model, we estimated the variance explained per phenotype variable for each gene. The “plotVarPart” function was used to plot the distribution of variance explained by different phenotypes and the “plotPercentBars” function was used to plot the top variable genes within each phenotype variable.

### Analysis of public snRNA-seq data of epilepsy and non-epilepsy human brain tissue

Gene counts and metadata of human cortex snRNA-seq data were obtained from the CELLxGENE database (https://cellxgene.cziscience.com/collections/ 35928d1c-36fc-4f93-9a8d-0b921ab41745). L2/3 IT neurons (79,631) were extracted to perform downstream analysis using Seurat R package. Briefly, top 2,000 highly variable genes selected by Variance Stabilizing Transformation (VST) were used to perform PCA analysis. Further harmony function was used to reduce the batch effect across different donors. Top 20 PCs were used to perform UMAP projection and clustering analysis. Similar to the previous section, GSVA scores of PY2 subtype signatures were calculated and mapped to L2/3 IT neurons of snRNA-seq data. Based on the distribution of PY2 GSVA scores across L2/3 IT neurons, those neurons with PY2 GSVA score > 0.6 were defined as PY2-like cells. Fisher's exact test was used to evaluate the proportion of PY2-like cells between epilepsy and non-epilepsy neurons. We further performed the comparison of PY2 GSVA scores between epilepsy and non-epilepsy PY2-like neurons using Wilcoxon test.

### EASI-FISH and imaging of thick brain slice after patch-clamp recordings

Following whole-cell patch-clamp recordings, human brain slices were post-fixed in 4% paraformaldehyde (PFA) in 1x PBS for 10-12 hours. Subsequently, slices were washed twice with 1x PBS and stored in 70% ethanol for up to 3 weeks. Before standard EASI-FISH processing, the slices were rehydrated in 1x PBS and incubated overnight with 1:1000 Alexa Fluor 555-conjugated streptavidin diluted in 1x PBS with 0.1% Tween-20 (PBST). Following incubation, the slices were rinsed twice with 1x PBS for 20 minutes each. The slices were then mounted onto glass-bottom dishes

(Cellvis) and imaged using an FV3000 confocal microscopy (Olympus) equipped with a 20x/1.0 objective. Regions of interest (ROIs) were identified and imaged. After image acquisition, the brain tissue was dissected under an MVX10 stereo microscopy (Olympus). ROIs were carefully dissected into smaller pieces (1-4 $mm^2$) containing the recorded cells.

These dissected tissue samples were then processed using a modified version of the previously described EASI-FISH protocol (6). Briefly, the small dissected tissues underwent RNA anchoring, gelation, Proteinase K digestion and DNase I digestion. Subsequently, 3 rounds of HCR RNA-FISH were performed to detect the expressions of 6 candidate marker genes (*NFKBIA, CDKN1A, NEFM, CCL2, CUX2, NEFH*), with 2 genes in each round, because the 546-channel was occupied by streptavidin. Prior to imaging, the samples were stained with 5 μg/ml 4',6-diamidino-2-phenylindole (DAPI) for 2 x 30 minutes. All samples were then imaged in PBS with 2x expansion using a Zeiss Lightsheet 7 microscope. Imaging was conducted using a 20x water-immersion objective (20x/1.0 W Plan-Apochromat Corr DIC M27 75 mm, RI=1.33) with 1x zoom. Images were collected at a pixel resolution of 0.23 x 0.23 μm (post-expansion) and a z-step size of 0.4 μm with single camera detection across four tracks: the 405 nm, 488 nm, 546 nm and 669 nm channels. After image acquisition, the probes and HCR hairpins were removed using DNase I.

**EASI-FISH data processing**

All 3D FISH image stacks were stitched using the Fiji Stitching Plugin (7). The stitched 3D image volumes from each round were then registered based on their DAPI signals using a customized Matlab script (8). To generate 3D cell boundaries, 3D cell segmentation was performed on the DAPI signals with Cellpose 3.0 (9) using a customized model, which was optimized for the human brain samples. The 3D segmentation results were further manually inspected and corrected. mRNA transcript signal was identified with a customized Fiji macro utilizing a RS-FISH (10) algorithm, which robustly identifies single-molecular spots in 3D with high precision. A total of 16372 cells were identified from 7 slice of 4 patient. For subsequent analysis, only cells expressing CUX2, NEFM and NEFH genes were selected, resulting in a refined dataset of 7688 cells (approximately 47% of the initial population, PY1 cluster: 5319 cells, PY2 cluster: 2369 cells). The cell-by-gene expression matrix was normalized using z-score. Unsupervised hierarchy clustering was employed to classify these neurons based on their expression profiles of six genes (*NFKBIA, CDKN1A, CCL2, NEFM, CUX2, NEFH*). Correlation distance and average linkage were utilized as the distance metric and linkage method, respectively, to group cells with similar expression patterns. The hierarchical clustering results were visualized using a two-dimensional UMAP (Uniform Manifold Approximation and Projection) embedding for dimensionality reduction and improved representation of the high-dimensional gene expression data in a lower-dimensional space. To assess the relationship between gene expression and cell size in the PY1 and PY2 clusters, we measured both slope and density. The slope was determined by performing a linear fit ($f(x) = k*x + b$) for expression (spot counts) of each gene against the corresponding volume ($\mu m^3$) of individual cells within the PY1 and PY2 clusters. The density of each recorded cell was calculated by dividing its gene expression (spot counts) by its respective volume ($\mu m^3$).

**Immunohistochemistry staining**

Surgical specimens were cut into slabs and immediately fixed in 4% paraformaldehyde and dehydration with 30% sucrose solutions for 24-48 h. Brain cryosections (10 - 20 μm thickness) were prepared from the brain tissue slabs using cryomicrotome

(Thermo Scientific, Cyrostar NX50) and then stored at -80°C. Before immunostaining experiments, slice sections were dried at 60°C for 10 min and rehydrated with 0.01M PBS for 3 times, 5 min each. Slice sections were then immersed in sodium citrate buffer (containing sodium citrate 9 mM and citric acid 1 mM) for 25 min for antigen retrieval. After air-cooled to room temperature, slice sections were washed with 0.01M PBS and then incubated with 10% donkey serum blocking buffer (10% donkey serum with 0.1% Triton X-100 dissolved in 0.01M PBS) for one hour to block non-specific antibody binding. The sections were incubated overnight at 4°C with primary antibodies, listed as follows: NeuN (chicken, 1:500, Millipore, ABN91), SMI311 (mouse, 1:500, BioLend, 837801), phosph-S6 (rabbit, 1:500, Cell signaling technology, 5364), P21 (rabbit, 1:200, Abcam, ab188224), COX2 (rabbit, 1:200, Abcam, ab15191), CCL2 (rabbit, 1:200, Sigma, HPA019163), P53 (rabbit, 1:200, Sigma, HPA051244), γ-H2AX (H2AYX, rabbit, 1:200, Abcam, ab11174), NFKBIA (rabbit, 1:200, Sigma, HPA029207), Lamin B1 (rabbit, 1:200, Abcam, ab16048). After a series of washes in PBS-T (with 0.1% Triton X-100, 3 times, 30 min each time), brain sections were incubated with the following fluorescein-conjugated secondary antibodies for one hour at room temperature, donkey anti-mouse Alexa Fluor 647 (1:1000, Invitrogen A31571), donkey anti-chicken Alexa Fluor 488 (1:1000, Jackson 703-545-155) and donkey anti-Rabbit Alexa Fluor 555 (1:1000, Thermofisher A31572). Secondary antibodies were removed and the slice sections were washed with PBS-T thoroughly (4 times, 30 min each time). To reduce the lipofuscin autofluorescence, the sections were washed in situ with freshly-prepared 70% alcohol for 5 min, incubated with 0.3% Sudan black (0.3 g in 100 mL 70% alcohol) for 15 min, then washed 3 times in 0.1 mol/L PBS-T for 10 min each. Cellular nuclei were stained with 4,6-diamidino-2-phenylindole (DAPI) for 2 min and then washed with 0.01M PBS. Finally slice sections were mounted with glass with FluorSave$^{TM}$ Reagent (Millipore, 345789) and imaged with confocal microscope (Olympus, FV3000).

β-galactosidase (SA-β-Gal) staining experiment was carried out based on senescence β-galactosidase staining kit (Cell signaling technology, 9860). Before β-Gal staining experiments, slice sections were dried at 60°C for 10 min and rehydrated with 0.01M PBS for 3 times, 5 min each. Post-fixation was done by addition of 1X Fixative Solution for 15 min at room temperature. Slice sections were washed with 0.01M PBS and then incubated overnight at 37°C in a dry incubator with the β-galactosidase staining solution. Then β-galactosidase staining solution was removed and washed 3 times, 10 min each, and the slice sections were mounted with FluorSave$^{TM}$ Reagent (Millipore, 345789) for imaging with an upright fluorescent microscope (Nikon ECLIPSE Ni).

**Electroencephalogram (EEG) recordings and seizure detection**

Mice were housed and treated according to the guidelines of the Zhejiang University (ZJU) Laboratory Animal Care and Use Committee and were approved by the Animal Advisory Committee at ZJU (ZJU20210231). CamKII-cre mice (The Jackson Laboratory, 005359) were crossed with *Pten*$^{flox/flox}$ mice (The Jackson Laboratory, 006440) to generate CamKII-cre::*Pten*$^{-/-}$ mice. CamKII-*Pten*$^{-/-}$ or *Pten*$^{flox/flox}$ control mice (6-10 postnatal weeks) were anesthetized with isoflurane and head fixed with a stereotaxic frame (RWD Life Sciences). Custom-made three-channel EEG electrodes were implanted to the skull of mice. The recording electrodes were placed in the frontal brain region (2.9 mm posterior and 3.2 mm lateral to bregma). The reference and ground electrodes were placed in the cerebellum region. The wires were connected with screws and secured with dental cement. A week after the surgery, 24 hours Video-EEG recordings were collected on a PowerLab 4 data acquisition unit (35PL3504/P, AD.) at 1KHz sample rate and analyzed offline with low-pass filter at 50 Hz using LabChart 8 (AD.).

The same littermates of CamKII::*Pten*$^{-/-}$ mice were randomly divided into rapamycin treatment group and control group (injected with corn oil only). Chronic rapamycin (Cat. #R-5000, LC laboratories. 10 mg/kg dissolved in corn oil) treatment was intraperitoneally (i.p.) injected to CamKII::*Pten*$^{-/-}$ mice for 10 days starting from 4 postnatal weeks (5 consecutive days from Monday to Friday, for 2 weeks) and continued with 2nd trial of 10 days treatment starting from 8 postnatal weeks. After rapamycin treatment, the CamKII::*Pten*$^{-/-}$ mice were monitored with 24-hour video-EEG recordings for 5 consecutive days and euthanized at 12 postnatal weeks. CamKII::*Pten*$^{-/-}$ mice are harvested following the detection of status epilepticus that is determined by the detection of repeated tonic-clonic seizures and lasts more than 5 minutes. Mice were euthanized with 1% sodium pentobarbital (50 mg/kg, i.p.) and perfused with 4% PFA, subsequently the mice brain were harvested and sectioned with a cryomicrotome for immunohistochemistry staining described above.

**Intra-hippocampal kainic acid (KA) injection induced acute and chronic epilepsy mouse model**

Mice were anesthetized with 1% sodium pentobarbital and were placed on a stereotaxic frame (RWD Life Science, 68046), where anesthesia was maintained with 1% - 1.5% isoflurane throughout the surgery. Before stereotaxic injection, the mice hair was clipped, the skull was exposed and the surgical area was sterilized with iodine povidone. Kainic acid (KA: 0.5 mg/ml in PBS. Abcam, ab120100) or PBS were slowly injected (0.4 μl over 5 min) with a manual syringe pump into the right hippocampus (coordinates: Y=2.0 mm, X=1.3 mm, Z=1.6 mm) by a thin glass capillary with a tip diameter of 50-100 μm. The needle was placed for additional 2 min after injection to prevent reflux along the injection trajectory and was then slowly lifted. The mice were placed on a heating pad to recover from the surgery and monitored for seizure in homecage. After injection with KA, all mice developed acute epileptic seizures, 20 ~ 30 % of mice died from status epilepticus (SE) shortly after the surgery. Three days after KA induced acute seizures, mice were euthanized and the mice brain were harvested for immunohistochemistry analysis.

Two months after KA injection, the mice were implanted with custom-made three-channel EEG electrodes in the skull and were monitored with 24-hour video-EEG recordings for 5 consecutive days following procedures described above. To record focal seizures, the recording electrode were implanted to the hippocampus ipsilateral to the KA injection site (coordinates: Y=2.9mm，X=3.2mm，Z=3.2mm). Electrographic spikes that had frequency higher than 2 Hz and amplitude higher than 3 times baseline with a minimal duration of 10 s were identified as rhythmic epileptiform activity. Bursts of high-voltage sharp waves with no obvious convulsive behaviors were identified as non-convulsive focal seizures (FS). Seizure events that are more than 10 s interval were considered as separate events. Paroxysmal EEG activity with post seizure depression along with obvious convulsive behaviors were identified as generalized tonic-clonic seizures (GS). After EEG recording, the mice were euthanized and the mice brain were harvested for immunohistochemistry analysis.

**Imaging processing, cell segmentation and colocalization analysis**

Fluorescent signals were acquired using a confocal laser scanning microscope (Olympus, FLUOVIEW IX83-FV3000). Single plane confocal images were acquired using 20X (0.75 NA, pixel size: 0.621 μm/pixel) and 40X (1.30 NA, pixel size: 0.311 μm/pixel) dry immersion objective. The co-localization of fluorescent signals from different marker genes was analyzed with Fiji software (ImageJ, National Institutes of Health). Cell number was counted from confocal images in a region of interest (ROI, 636.4 × 636.4 $\mu m^2$) using ImageJ and Imaris 9.7.1 software (Oxford Instruments Group). Each marker gene expression was quantified from confocal images of at least three individual brain samples of drug-resistant epilepsy patients with different pathologies (FCD: focal cortical dysplasia; TLE: temporal lobe epilepsy; PT: peri-tumor cortex of patients with tumor induced epilepsy) as well as control brain samples with no history of epilepsy (Control). The number of β-Gal positive cells was calculated in images with ROI (1204 x 920 $\mu m^2$) from individual brain samples. The immunostaining protocol and image acquisition parameters of the same antibody were kept consistent across different brain samples.

We performed the imaging processing analysis using Cellpose Python package(11), Fiji (ImageJ, National Institutes of Health) and customized Matlab (MathWorks) scripts. Briefly, Cellpose was used to identify the cell candidates; to reduce false positive in low signal-noise-ratio images, foreground cell masks were generated using Fiji thresholding; finally, a Matlab script was used to determine the masks of individual cells by combining the cell candidates and foreground cell masks. In detail, first, raw images were pre-processed by a customized Fiji macro. Each image was split and saved as two images: one merging the DAPI and interest marker channels for segmenting cell candidates with Cellpose; the other one containing only the interest marker channel for generating foreground cell mask with Fiji. After the image pre-processing, we segmented the merged images into candidate cells using Cellpose with different diameter parameter based on their mean cell diameter in “cyto2” mode. Next, we generated foreground cell masks from the single marker images using a customized Fiji macro. Images were “despeckle” to remove non-specific signals, gaussian blurred to reduce the background noise and automatically thresholded to generate the foreground cell masks in the single marker images. Due to the large signal variations of the stained markers, we used “Ostu” algorithm to threshold NeuN channel and applied “Triangle” or “Yen” thresholding algorithm for other marker channels to optimize the background-foreground segmentation. In addition, we used an “Area open” filter to remove the foreground cell masks with an area smaller than 60 pixels (Lamin B1), 100 pixels (CCL2, COX2, γH2AX, NFKBIA, P21, P53) or 150 pixels (NeuN). Subsequently, we used a customized Matlab script to determine individual cells. If a cell candidate had > 60 pixels in the nucleus mask and > 60, 100 or 150 pixels in the foreground cell mask, we assigned it as a real cell. Otherwise, we removed it from the candidate cell list. These real cells were automatically counted with a customized Fiji macro. After generating the real cell masks for each channel, we intersected the real cell masks between the NeuN image and the interest marker image as the colocalization cell masks. To avoid intersecting two closely adjacent cells, the colocalization cell masks with an area < 60 pixels were removed. Colocalization masks were also automatically counted by the same Fiji macro described above.

### Supplemental Figures and Figure legends 1-10

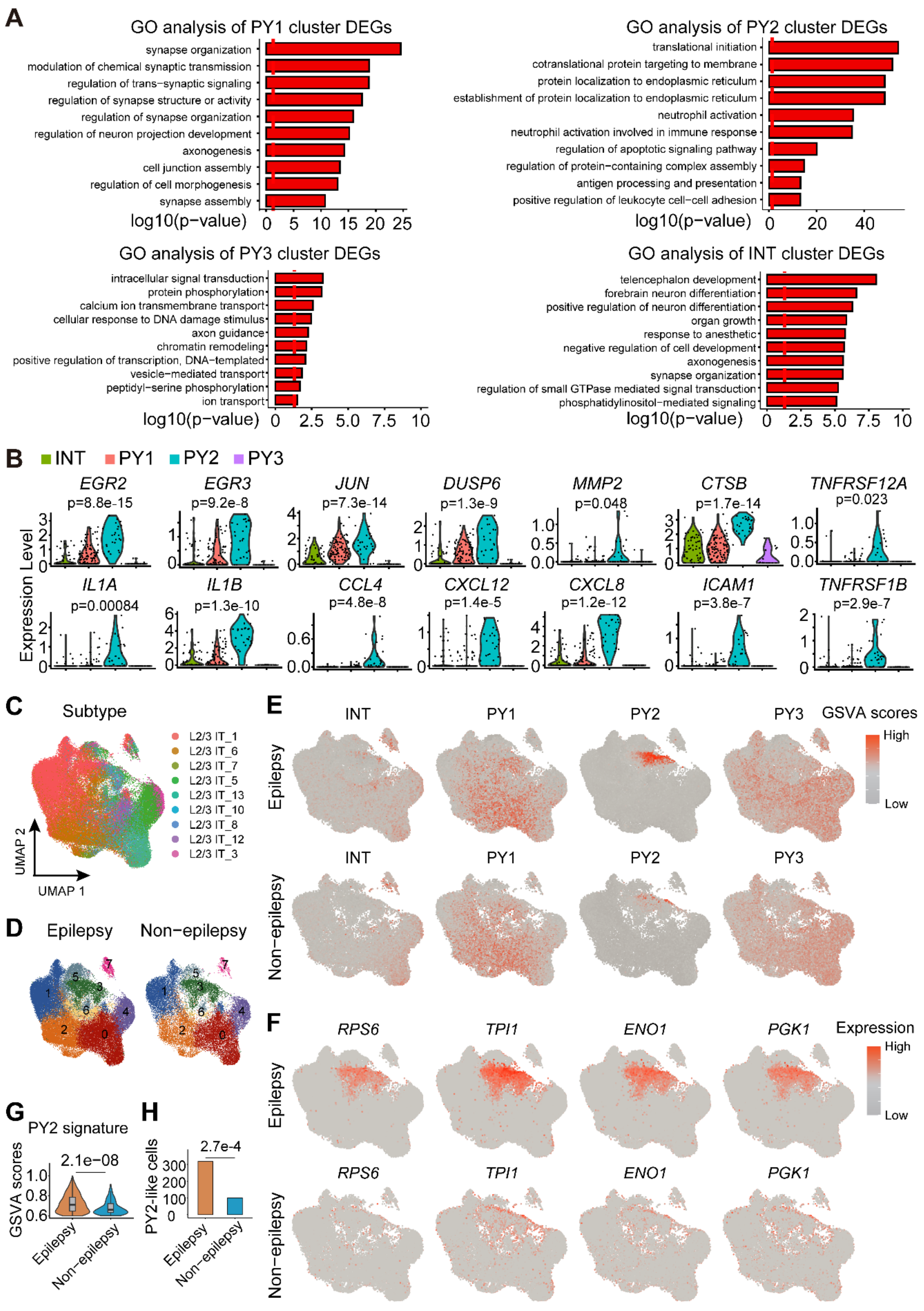

**Supplemental Figure 1. Differential gene expression analysis of distinct transcriptomic neuron subclasses. Related to Figure 1.** **(A)** Gene ontology analysis (GO) of differential expressed genes (DEGs) in INT interneuron, PY1, PY2 and PY3 pyramidal neuron clusters. **(B)** Violin plots showed representative DEGs in PY1 and PY2 pyramidal neuron clusters. Y axis indicated relative expression level. **(C)** UMAP representation of 79,631 L2-3 IT neuron subtypes of adult human brain from public snRNA-seq dataset indicate the enrichment of PY2-like cells in epilepsy patients, compared with non-epilepsy controls. **(D)** Unsupervised re-clustering of L2-3 IT neurons from brain samples of epilepsy or non-epilepsy groups after Harmony integration. **(E)** Feature plots displaying the comparison of GSVA scores of four patch-seq neuron subtype signatures (INT, PY1, PY2 and PY3), between epilepsy patients (top) and non-epilepsy controls (bottom). **(F)** Feature plots showing expression patterns of four mTOR pathway genes (RPS6, TPI1, ENO1, PGK1) enriched in PY2-like neurons in brain samples from epilepsy patients (top), compared with L2-3 IT neurons non-epilepsy control brain samples (bottom). **(G)** Violin plots showing the patterns of PY2 GSVA scores in the PY2-like cluster 5 between epilepsy patients and non-epilepsy controls. **(H)** Bar plots showing the number of PY2-like cells in different disease states. Fisher's exact test, *P* = 0.00027, odds ratio = 1.5.

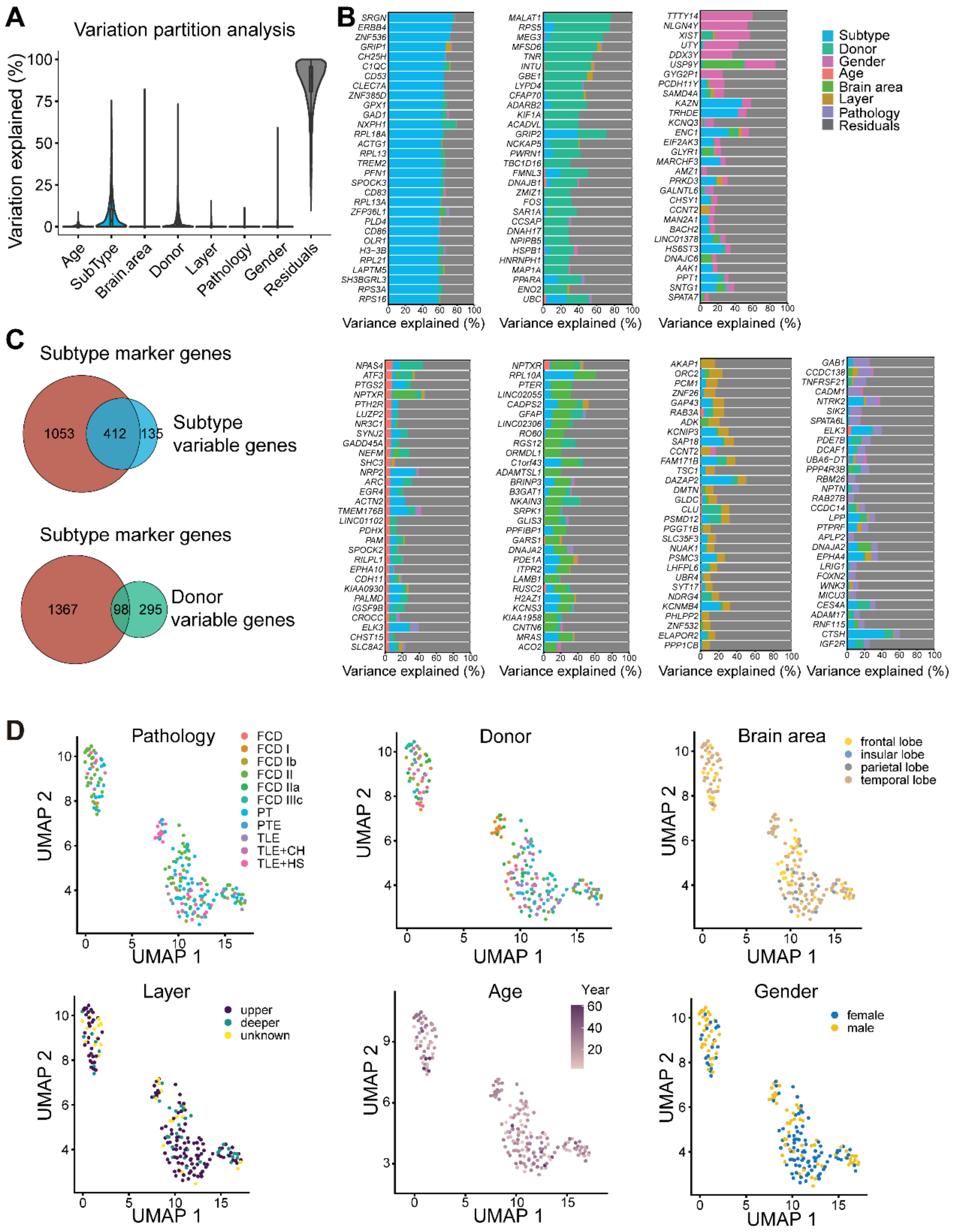

**Supplemental Figure 2. Variation partition analysis reveals interindividual variation of gene expression associated with biological factors. Related to Figure 1.** **(A)** Violin plots to show the percentage of gene variance explained by biological factors including age, subtype, brain area, donor, layer, disease pathology and gender (seven covariates). **(B)** Bar plots to present top 30 genes showing high variability within subtypes, donor, gender, age, brain area, layer and pathology. **(C)** Venn diagram showing the shared genes between subtype-specific marker genes and highly variable genes within subtypes (upper panel) or highly variable genes within donor (lower panel). **(D)** Uniform Manifold Approximation and Projection (UMAP) plots showing a lack of correspondence between transcriptomic cell subtypes with pathology, donor, brain area, cortical layer distribution, age and gender.

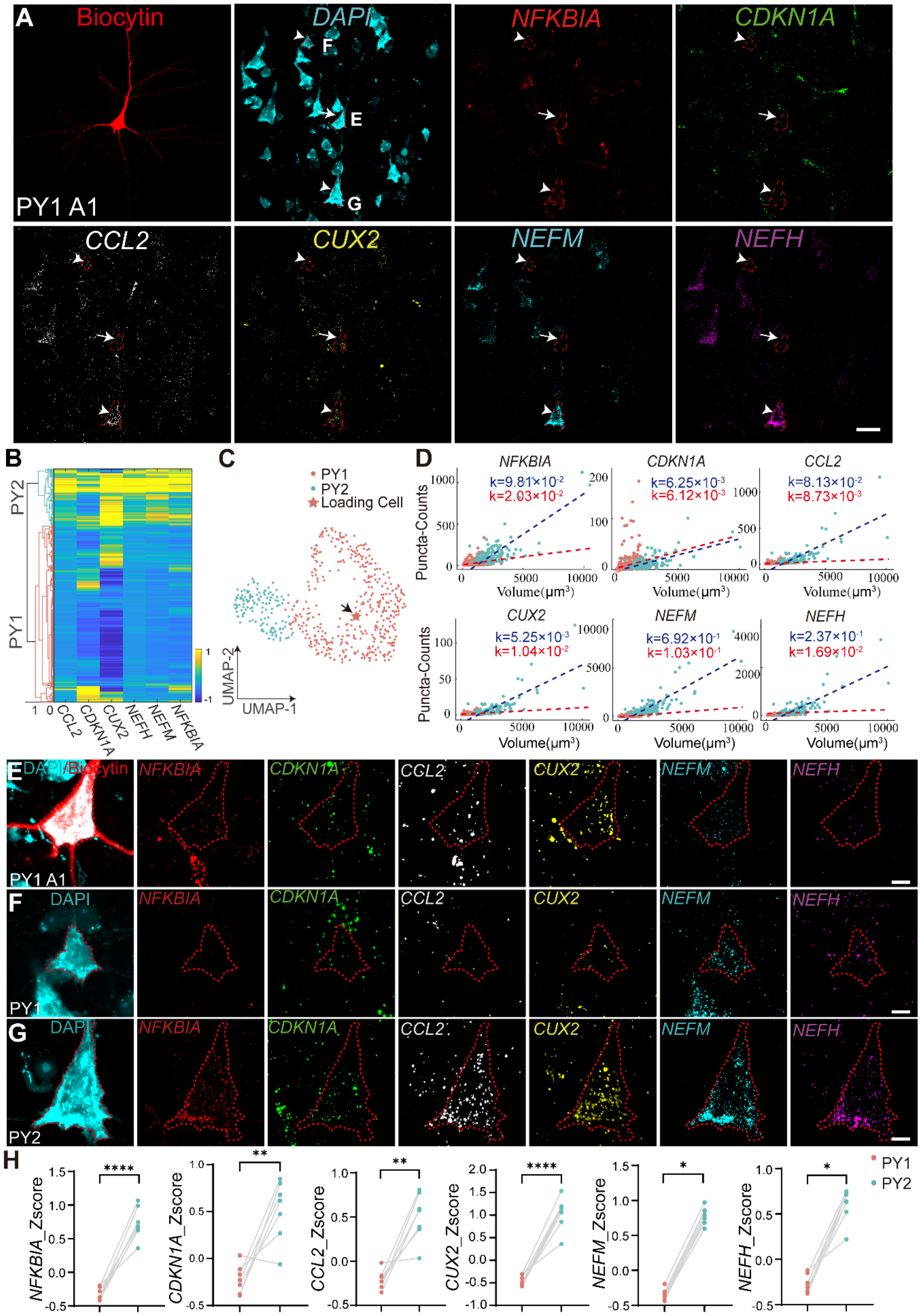
A
Biocytin
DAPI
NFKBIA
CDKN1A
PY1 A1
CCL2
CUX2
NEFM
NEFH
B
PY2
PY1
C
PY1
PY2
Loading Cell
UMAP-2
UMAP-1
D
Puncta-Counts
Volume(μm³)
E
F
G
H
PY1
PY2

**Supplemental Figure 3. Multiplex EASI-FISH reveals molecularly-defined PY1 neurons and their electrophysiological and morphological characteristics. Related to Figure 3. (A)** Example FISH images showing the expression of *NFKBIA, CDKN1A, CCL2, CUX2, NEFM, NEFH* in a ROI with biocytin loaded PY1 neuron in a brain slice from human drug-resistant epilepsy. Scale bar, 50 μm. **(B)** Heatmap of six differentially expressed genes (DEGs) of two molecularly-defined clusters in an example brain slice. **(C)** Uniform Manifold Approximation and Projection (UMAP) analysis showing distribution of PY1 and PY2 neurons in the same brain slice as in B. **(D)** Scatter plots showing the gene expression densities of *NFKBIA, CDKN1A, CCL2, CUX2, NEFM* and *NEFH* in PY1 and PY2 neuron clusters in the same brain slice as in B and C. Red dashed line, PY1 cluster density fitting line; Blue dashed line, PY2 cluster density fitting line. **(E-G)** Representative images showing the expression of candidate marker genes in biocytin-labeled PY1 neuron (E), molecular-defined PY1(F) and PY2(G) pyramidal neurons. **(H)** Statistical results showing the mRNA counts in PY1 and PY2 neurons (n = 7688 neurons, 7 brain slices from 4 individual samples). *NFKBIA*, $P < 0.0001$; *CDKNIA*, $P = 0.0048$; *CCL2*, $P = 0.0016$; *CUX2*, $P < 0.0001$; *NEFM*, $P = 0.0156$, *NEFH*, $P = 0.0156$; Each data point indicates the mean value from a sample; gray lines connect data points from the same sample. *$P < 0.05$; ** $P < 0.01$; ***$P < 0.001$; ****$P < 0.0001$. n.s. non-significant. Normality and Lognormality Tests were used to determine whether the data fit the normal distribution, two-tailed paired t-test is employed when the data follows a normal distribution, whereas the non-parametric two-tailed Wilcoxon signed rank test is utilized in cases where the data does not conform to a normal distribution.

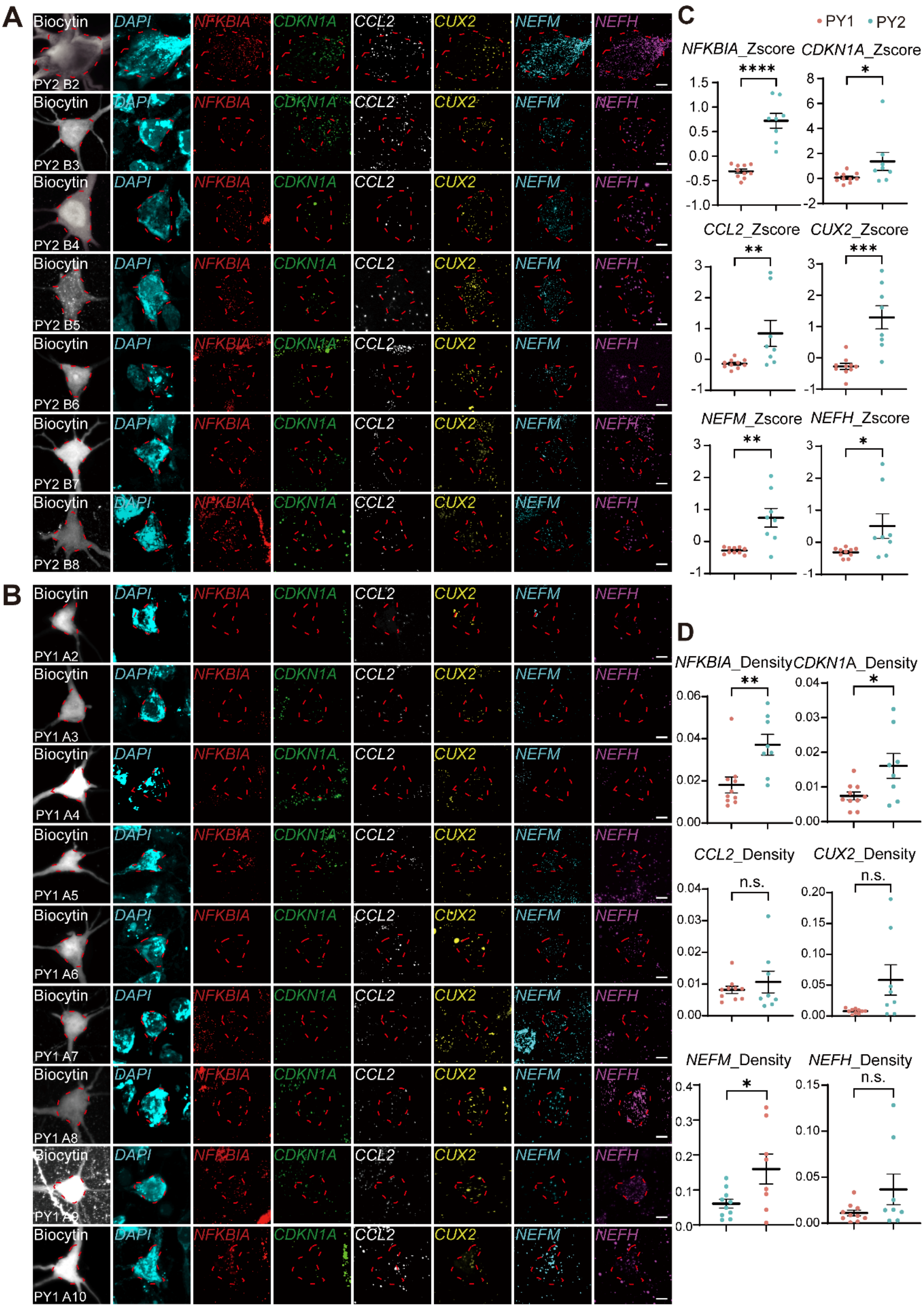
A
B
C
D
Biocytin
DAPI
NFKBIA
CDKN1A
CCL2
CUX2
NEFM
NEFH
PY2 B2
PY2 B3
PY2 B4
PY2 B5
PY2 B6
PY2 B7
PY2 B8
PY1 A2
PY1 A3
PY1 A4
PY1 A5
PY1 A6
PY1 A7
PY1 A8
PY1 A9
PY1 A10
PY1
PY2
NFKBIA_Zscore
CDKN1A_Zscore
CCL2_Zscore
CUX2_Zscore
NEFM_Zscore
NEFH_Zscore
NFKBIA_Density
CDKN1A_Density
CCL2_Density
CUX2_Density
NEFM_Density
NEFH_Density
n.s.

**Supplemental Figure 4. Expression of candidate marker genes in biocytin-loaded PY1 and PY2 neurons in brain slices from human drug-resistant epilepsy. Related to Figure 3. (A** and **B)** FISH images showing the expression of *CDKN1A, NFKBIA, CCL2, CUX2, NEFM, NEFH* in biocytin-loaded PY1 (A) and PY2 (B) pyramidal neurons. Scale bar, 10 μm. **(C)** Statistical quantification showing the expression of *CDKN1A, NFKBIA, CCL2, CUX2, NEFM, NEFH* in PY1 pyramidal neurons compared with PY2 pyramidal neurons. *NFKBI1*, P < 0.0001; *CDKN1A*, P = 0.0196; *CCL2*, P = 0.0196; *CUX2*, P = 0.0002; *NEFM*, P = 0.0012; *NEFH*, P = 0.0343. **(D)** Statistical analysis of the gene expression density (Counts/Volume) in PY1 and PY2 neurons. PY1, n = 8; PY2, n = 10. *NFKBIA*, *P* = 0.0062; *CDKNIA*, *P* = 0.0221; *CCL2*, *P* = 0.8286; *CUX2*, *P* = 0.0545; *NEFM*, *P* = 0.0267; *NEFH*, *P* = 0.2370. Data are presented as mean ± SEM. **P* < 0.05; ***P* < 0.01; ****P* < 0.001; *****P* < 0.0001. n.s. non-significant. Normality and Lognormality Tests were used to determine whether the data fit the normal distribution, two-tailed paired t-test is employed when the data follows a normal distribution, whereas the non-parametric two-tailed Wilcoxon signed rank test is utilized in cases where the data does not conform to a normal distribution.

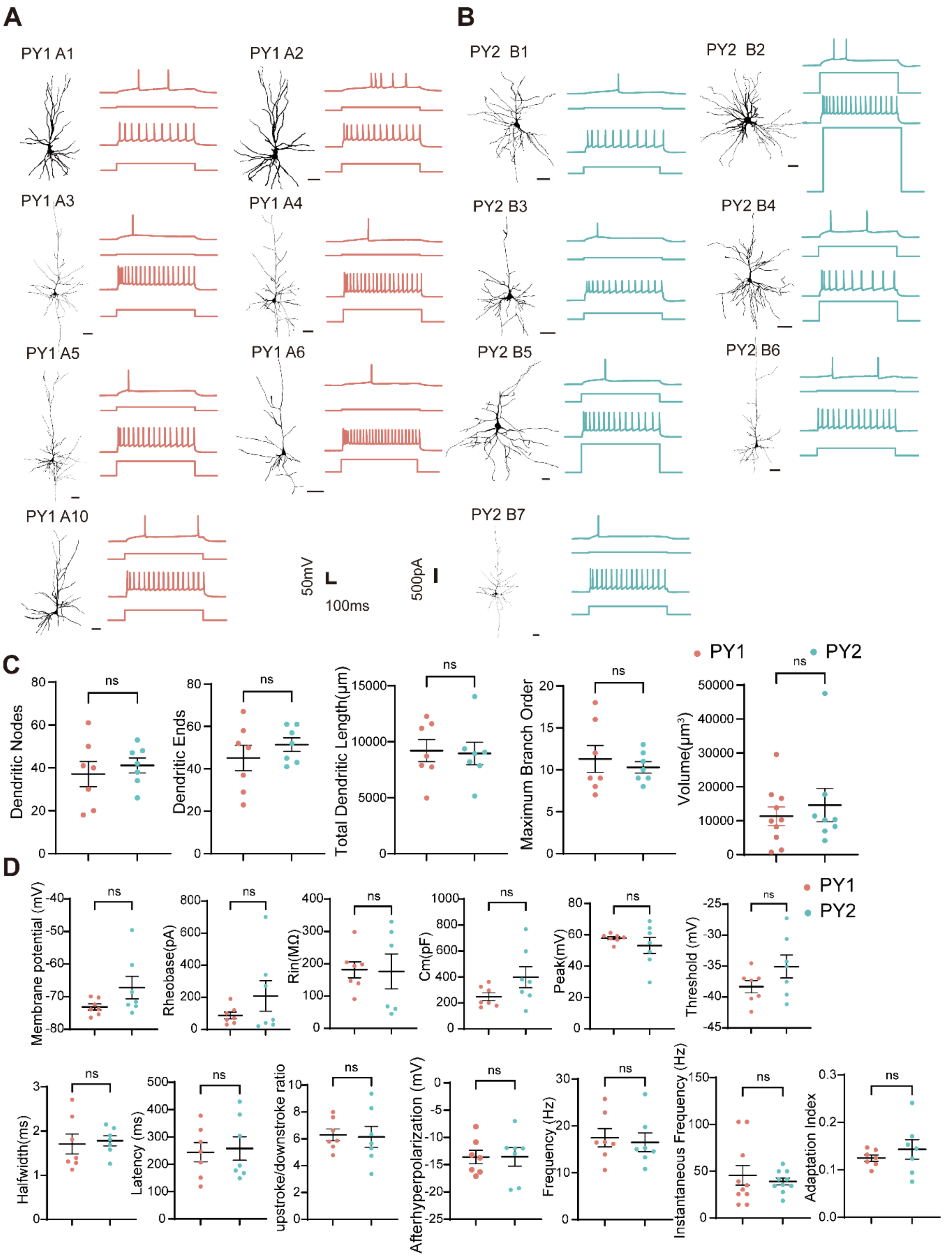


**Supplemental Figure 5. Electrophysiological and morphological characteristics of molecularly-defined PY1 and PY2 neurons. Related to Figure 3.** (**A** and **B**) Left, examples of action potential firing patterns of molecularly-defined PY1 (A) and PY2 neurons (B). Scale bars: 50 mV, 100 ms in AP firing patterns trace; 500 pA, in stimulation trace. Right, Reconstructed morphologies of PY1 (A) and PY2 (B) pyramidal neurons revealed by intracellular loading with biocytin. Scale bar, 100 μm. **(C)** Statistical analysis of morphological parameters and volume between PY1 and PY2 neurons. PY1, n = 7; PY2, n = 7; Dendritic nodes, $P$ = 0.5771; Dendritic ends, $P$ = 0.1882; Dendritic length, $P$ = 0.8618; Maximum branch order, $P$ = 0.5757; Volume, $P$ = 0.5492 (PY1, n = 10, PY2, n = 8); **(D)** Statistical analysis of electrophysiological parameters between PY1 and PY2 neurons. PY1, n = 7; PY2, n = 7; Membrane potential, $P$ = 0.1208; Rheobase, $P$ = 0.9225; Rin, $P$ = 0.9276; Cm, $P$ = 0.1069; Peak, $P$ = 0.3829; Threshold, $P$ = 0.1487; Halfwidth, $P$ = 0.7842; Latency, $P$ = 0.8079; upstroke/downstroke ratio, $P$ = 0.8717; Afterhyperpolarization, $P$ = 0.9837; Frequency $P$ = 0.7330; Instantaneous Frequency, $P$ = 0.6163; Adaptation Index, $P$ = 0.4075. n.s. non-significant. Two-tailed t-test.

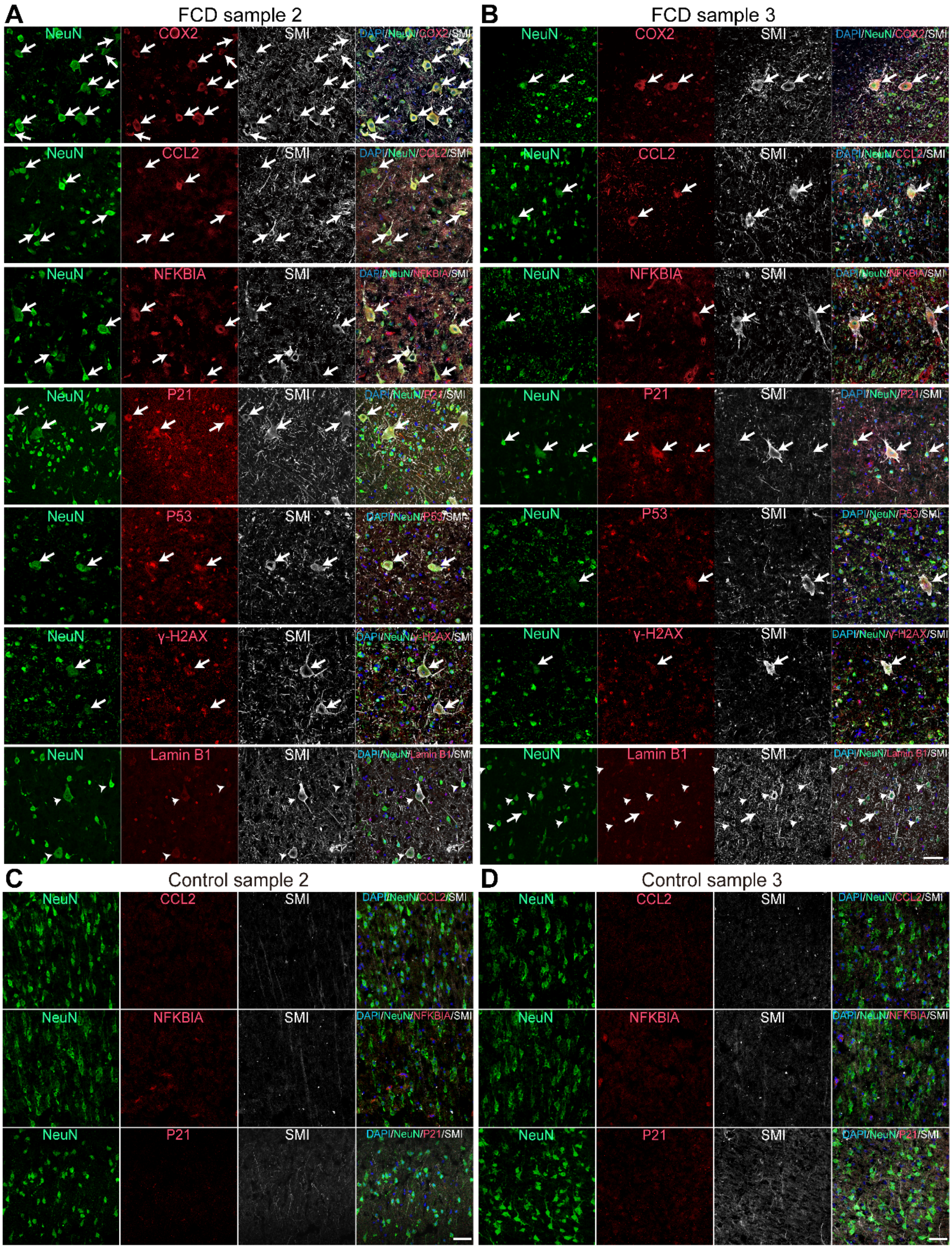
A
FCD sample 2
B
FCD sample 3
NeuN
COX2
SMI
DAPI/NeuN/COX2/SMI
CCL2
DAPI/NeuN/CCL2/SMI
NFKBIA
DAPI/NeuN/NFKBIA/SMI
P21
DAPI/NeuN/P21/SMI
P53
DAPI/NeuN/P53/SMI
γ-H2AX
DAPI/NeuN/γ-H2AX/SMI
Lamin B1
DAPI/NeuN/Lamin B1/SMI
C
Control sample 2
D
Control sample 3

**Supplemental Figure 6. Immunohistochemistry analyses of the expression of marker genes in dysmorphic neurons in two additional independent brain tissue samples of focal cortical dysplasia. Related to Figure 4 and Figure 5. (A** and **B)** Immunohistochemistry staining showing the expression of CCL2, NFKBIA, P21, P53, COX2, γ-H2AX and reduction of Lamin B1 in SMI-positive dysmorphic neurons in two independent brain samples of focal cortical dysplasia. **(C** and **D)** Immunohistochemistry staining showing the expression of CCL2, NFKBIA, P21 and SMI in two independent control brain samples with no history of epilepsy. Arrows indicated co-expression of PY2 DEG and senescent cell marker genes with SMI and NeuN. Arrowheads indicated no Lamin B1 expression were detected in SMI-positive histopathological neurons. Scale bar, 50 μm.

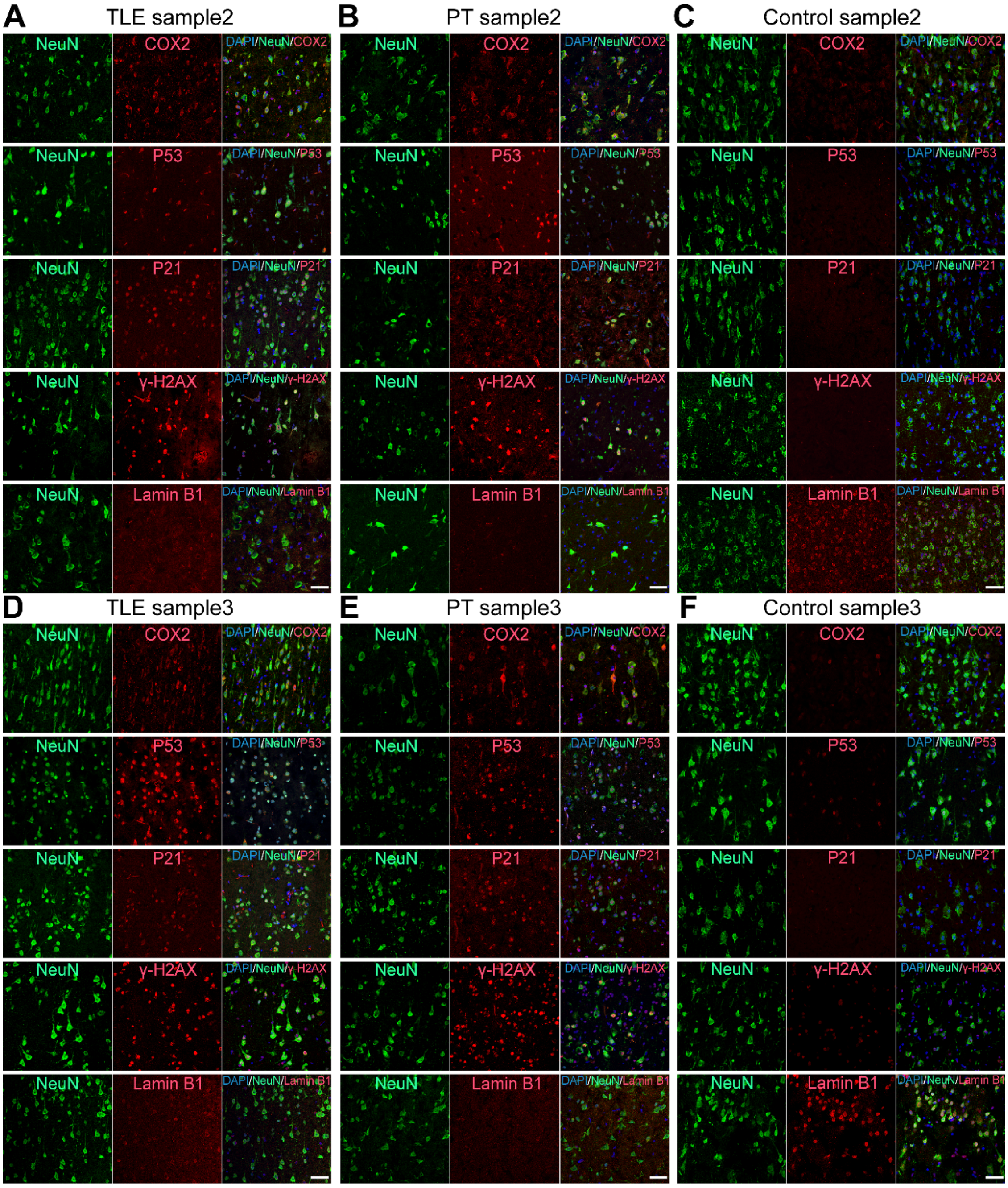


**Supplemental Figure 7. Expression of senescent cell markers in cortical neurons in the epileptic focus of additional brain samples from drug-resistant epilepsy patients with different pathologies. Related to Figure 6. (A** and **D)** Representative images showed the expression of senescent marker genes (COX2, P53, P21, γ-H2AX) and reduction of Lamin B1 in cortical neurons in the epileptic focus of TLE, PT (**B** and **E**), but not in control (**C** and **F**) brain samples. Scale bar, 50 μm.

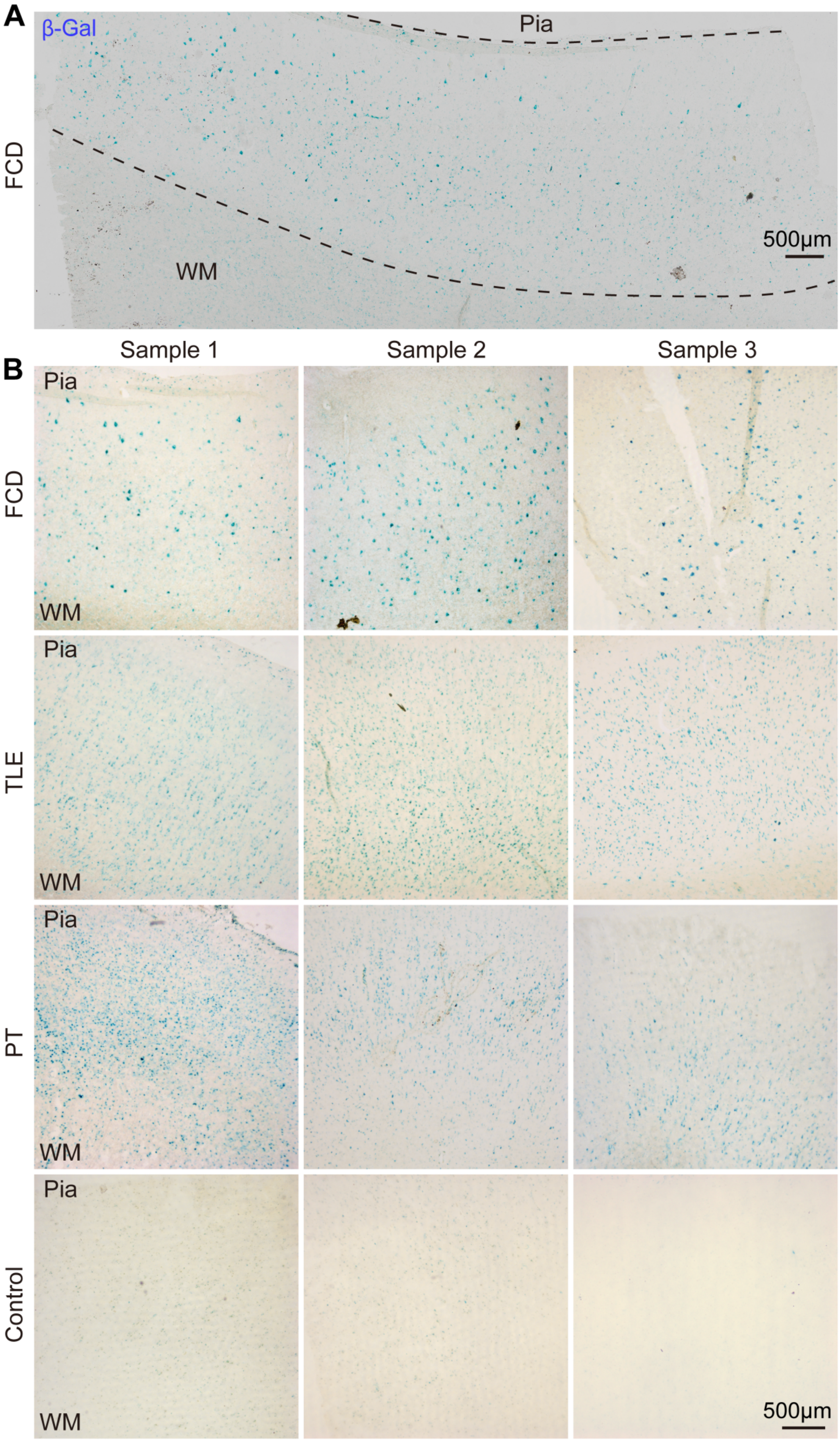
A
β-Gal
Pia
FCD
WM
500μm
Sample 1
Sample 2
Sample 3
B
Pia
FCD
WM
Pia
TLE
WM
Pia
PT
WM
Pia
Control
WM
500μm

**Supplemental Figure 8. Representative images showing the distribution of β-Gal positive cells in brain tissue from people with drug-resistant epilepsy, related to Figure 5. (A)** Representative tile-scan images showing β-Gal positive cells widely distributed in the brain tissue from people with drug-resistant epilepsy. **(B)** Representative images showing the wide distribution of β-Gal positive cells in the brain tissue from drug-resistant epilepsy with FCD, TLE, PT, but not in control cortical tissue with no history of epilepsy. N = 3 independent brain samples of each pathology.

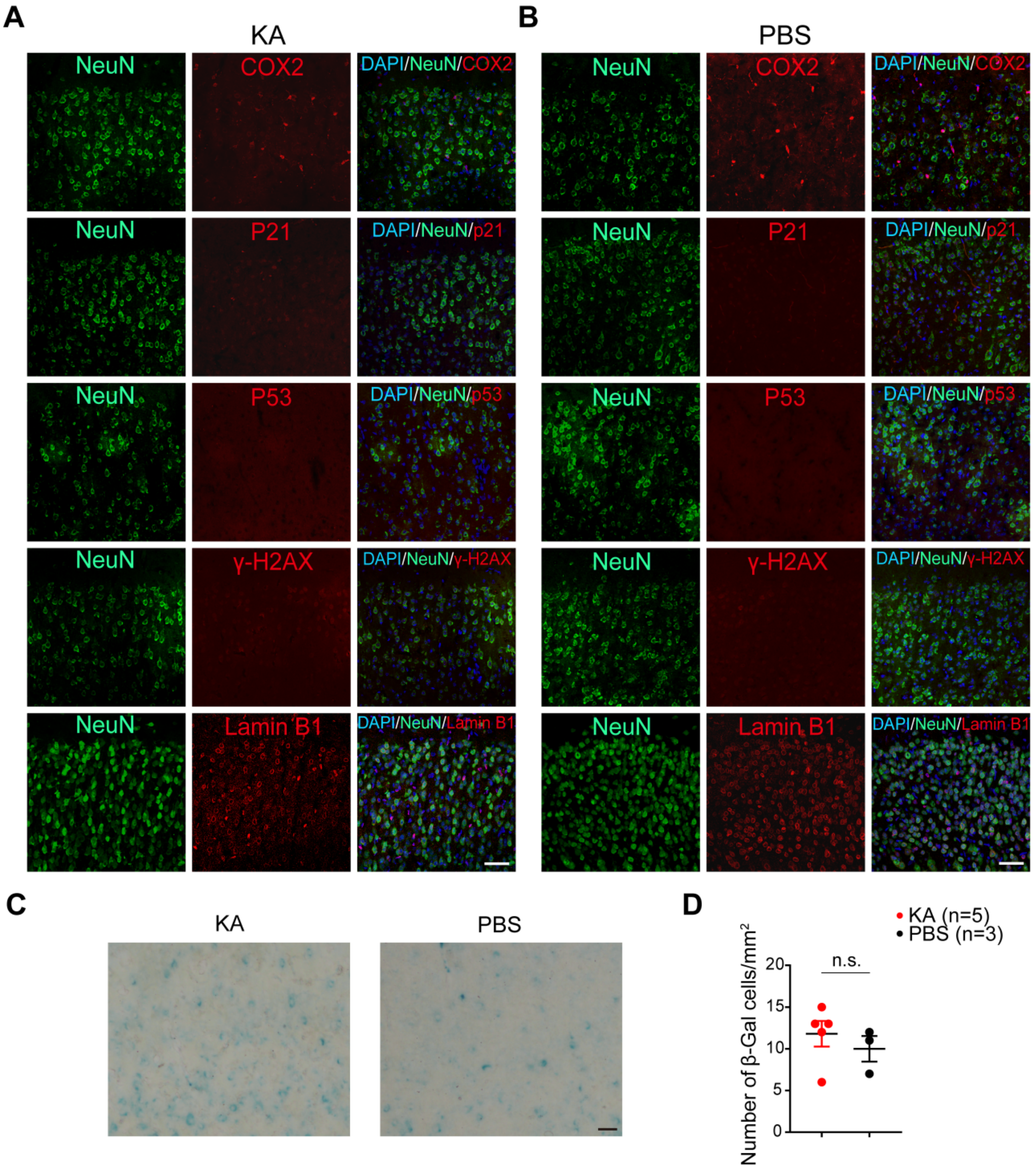


**Supplemental Figure 9. Acute epileptic activity did not induce the expression of senescent markers in cortical neurons of KA induced epilepsy mouse model. Related to Figure 7. (A-C)** Representative images showing the expression of senescent markers (COX2, P21, P53, γ-H2AX, Lamin B1) in the cortex of KA - treated (three days after KA injection) compared with PBS - treated mice (B). **(C)** Representative images and statistical results (**D**) showing the expression of SA-β-Gal in the cortex of KA - treated compared with PBS - treated mice. Data are presented as mean ± SEM. n.s. non-significant. Two-tailed Mann-Whitney test.

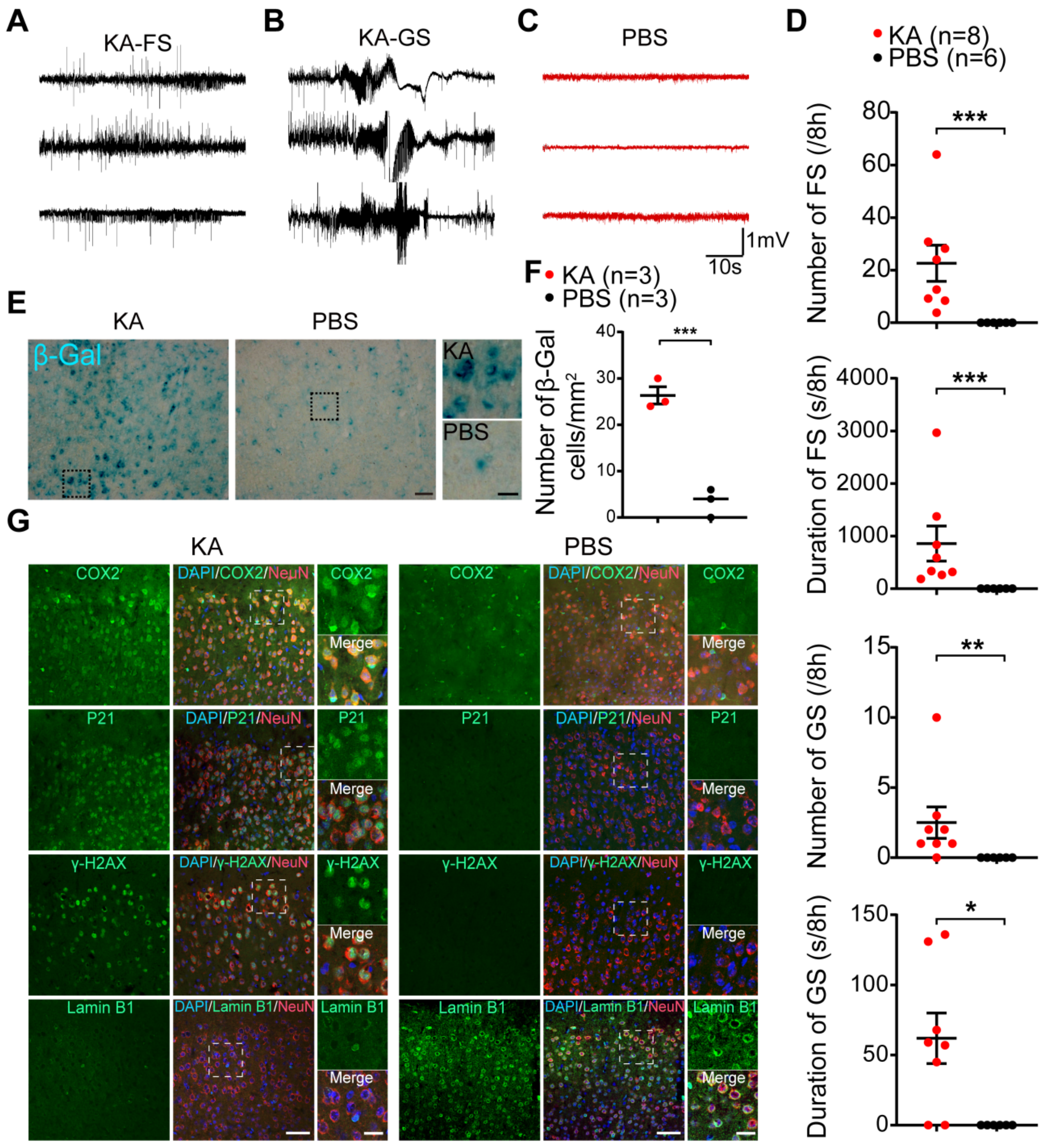


**Supplemental Figure 10. Neuronal senescent markers were induced by generalized tonic-clonic seizures in KA induced chronic epilepsy model. Related to Figure 7. (A-C)** Representative traces from EEG recordings for KA (black traces, B) and PBS treated (red traces, C) mice. **(D)** Statistical analysis of the average number of focal seizures (FS) or generalized tonic-clonic seizures (GS) per 8 hours in KA treated mice compared with PBS (control) treated mice. Number of FS (KA treated: 22.63 ± 6.88; PBS treated: 0.00 ± 0.00. *P* = 0.0007); Number of GS (KA treated: 2.50

± 1.12; PBS treated: 0.00 ± 0.00. *P* = 0.0033); Duration of FS (KA treated: 857.60 ± 331.60; PBS treated: 0.00 ± 0.00. *P* = 0.0007); Duration of GS (KA treated: 61.99 ± 18.07; PBS treated: 0.00 ± 0.00. *P* = 0.0150. Two-tailed *t*-test). **(E)** SA-β-Gal activity can be induced by spontaneous epilepsy in KA treated mice, but not PBS treated mice. Scale bar, 50 μm. **(F)** Statistical results showing the average number of SA-β-Gal positive neurons per 1 $mm^2$ field of view with 20× objective in (**E**). KA treated: 26.33 ± 1.86, n = 3; PBS treated: 3.33 ± 1.76, n = 3. P=0.0008. Two-tailed Mann-Whitney test. **(G)** Expression of senescence marker can be induced by epilepsy in KA treated mice, but not in PBS treated mice. The zoom-in images (right) show high magnification of co-expression of senescence markers with NeuN. Scale bar, 50 μm. Data are presented as mean ± SEM.

**Supplemental Table 1. Clinical data of the patients.**

| ID | Diagnosis | Epilepsy duration | Age at surgery | Monthly seizure frequency | Experimental objective | Brain Region |
|---|---|---|---|---|---|---|
| **FCD** | | | | | | |
| 1 | FCD IB | 3 | 6 | 150 | Patch-seq | L.F |
| 2 | FCD I | 6 | 9.5 | * | Patch-seq | R.F |
| 3 | FCD IIA | 1 | 1.5 | 30 | Immunofluorescence | L.F |
| 4 | FCD IIA | 3 | 6 | 30 | Immunofluorescence | L.T |
| 5 | FCD IIA | 1.5 | 2.08 | 30 | Patch-seq | L.F |
| 6 | FCD IIA | 4 | 24 | 1.5 | Patch-seq | R.F |
| 7 | FCD IIA | 28 | 33 | 75 | Patch-seq | R.F |
| 8 | FCD IIA | 20 | 43 | 30 | Patch-seq | L.F |
| 9 | FCD IIA | 12 | 20 | 20 | Patch-seq | L.F |
| 10 | FCD IIA | 5 | 13 | 195 | Patch-seq | L.F |
| 11 | FCD IIA | 3 | 6.08 | 30 | Patch-seq | L.T |
| 12 | FCD IIA | 8 | 10.33 | 10 | Patch-seq | L.F |
| 13 | FCD IIA | 20 | 32 | 4 | Immunofluorescence | L.P |
| 14 | FCD IIA | 3 | 4.25 | 30 | Immunofluorescence | R.F |
| 15 | FCD IIA | 4 | 13.25 | 2 | Immunofluorescence | L.F |
| 16 | FCD IIA | 10 | 22 | * | Immunofluorescence | L.F |
| 17 | FCD IIA | 12 | 28 | * | Immunofluorescence | R.O |
| 18 | FCD IIB | 17 | 26 | 105 | Immunofluorescence | R.F |
| 19 | FCD IIB | 1.5 | 2.58 | 600 | Immunofluorescence | L.F |
| 20 | FCD IIB | 8 | 12 | * | Patch-seq + staining | R.F |
| 21 | FCD IIB | 0.75 | 7.58 | 45 | Patch-seq | L.T |
| 22 | FCD IIB | 20 | 35 | 1 | Patch-seq | R.T |
| 23 | FCD IIB | 25 | 26 | 1.5 | Patch-seq | R.T |
| 24 | FCD IIB | 0.92 | 2.75 | * | Patch-seq + staining | L.P |
| 25 | FCD IIB | 10 | 13 | 90 | Patch-seq + staining | R.F |
| 26 | FCD IIB | 2 | 17 | 0 | Patch-seq | L.P |

| | | | | | | |
|---|---|---|---|---|---|---|
| 27 | FCD IIB | 5 | 16 | 45 | Immunofluorescence | L.F |
| 28 | FCD IIB | 10 | 13 | * | Immunofluorescence | R.F |
| 29 | FCD IIB | 2 | 2.75 | * | Immunofluorescence | R.F |
| 30 | FCD II | 4 | 48 | * | Patch-seq | L.T |
| 31 | FCD IIIC | 3 | 11.83 | * | Patch-seq | R.F |
| 32 | FCD I | 0.42 | 4 | * | Electrophysiology + FISH | L.F |
| 33 | FCD IIA | 9 | 16 | * | Electrophysiology + FISH | L.F |
| 34 | FCD I | 0.58 | 2 | * | Electrophysiology + FISH | L.F |
| **TLE** | | | | | | |
| 32 | Crypto | 10 | 28 | 3 | Immunofluorescence | R.T |
| 33 | Crypto | 10 | 34 | 0 | Immunofluorescence | R.T |
| 34 | Crypto | 10 | 22 | 15.3 | Patch-seq + staining | R.T |
| 35 | Crypto | 8 | 52 | 2.5 | Immunofluorescence | L.T |
| 36 | Crypto | 4 | 48 | 6 | Immunofluorescence | L.T |
| 37 | Crypto | 18 | 27 | * | Immunofluorescence | L.T |
| 38 | HS | 6 | 15 | 0.75 | Immunofluorescence | L.T |
| 39 | HS | 18 | 28 | 3 | Patch-seq | R.T |
| 40 | HS | 7 | 16 | 4 | Patch-seq + staining | L.T |
| 41 | HS | 26 | 31 | 135 | Patch-seq + staining | L.T |
| 42 | HS | 10 | 33 | 0.2 | Patch-seq | R.T |
| 43 | HS | 6 | 13.25 | 4.5 | Immunofluorescence | R.T |
| 44 | HS | 15 | 59 | 1.5 | Immunofluorescence | L.T |
| 45 | HS | 8 | 27 | * | Immunofluorescence | L.T |
| 46 | HS | 14 | 30 | 4.5 | Patch-seq | R.T |
| 47 | HS | 15 | 32 | * | Patch-seq | R.H |
| 48 | CH | 1 | 25 | 6 | Immunofluorescence | R.T |
| 49 | CH | 5 | 61 | 60 | Patch-seq | L.T |
| **PT** | | | | | | |
| 50 | GG | 0.25 | 29 | 3 | Immunofluorescence | R.T |

| | | | | | | |
|---|---|---|---|---|---|---|
| 51 | GG | 30 | 54 | * | Immunofluorescence | L.T |
| 52 | GG | 2 | 27 | 0.1 | Immunofluorescence | R.T |
| 53 | GG | 10 | 22 | 2.5 | Patch-seq + staining | R.T |
| 54 | GG | 0.33 | 30 | 3 | Patch-seq + staining | R.T |
| 55 | GG | 0.25 | 22 | 1.5 | Patch-seq + staining | L.T |
| 56 | GG | 0.67 | 2.42 | 1.5 | Patch-seq + staining | L.T |
| 57 | GG | 0.17 | 20 | * | Patch-seq | R.T |
| 58 | GG | 0.75 | 56 | * | Patch-seq | R.T |
| 59 | GG | 0.5 | 2 | * | Immunofluorescence | L.T |
| 60 | GG | 8 | 15 | * | Immunofluorescence | L.T |
| 61 | GG | 1.5 | 2 | * | Immunofluorescence | L.F |
| 62 | PXA | 1 | 21 | 0.8 | Patch-seq | R.T |
| 63 | PXA | 2 | 6.25 | 60 | Patch-seq + staining | R.T |
| 64 | PXA | 5 | 41 | 1 | Immunofluorescence | R.T |
| 65 | PA | 0.75 | 18 | 35 | Immunofluorescence | R.T |
| 66 | PA | 8 | 32 | 2 | Immunofluorescence | R.F |
| 67 | PLNTY | 6 | 11 | * | Electrophysiology + FISH | R.P |
| **Traumatic epilepsy** | | | | | | |
| 67 | TE | 20 | 30 | 2 | Patch-seq | R.P |
| **Control** | | Age at death | | | | |
| 68 | Control | 0.5 | / | / | Immunofluorescence | R.F |
| 69 | Control | 50 | / | / | Immunofluorescence | R.F |
| 70 | Control | 51 | / | / | Immunofluorescence | R.F |
| 71 | Control | 42 | / | / | Immunofluorescence | R.F |
| 72 | Control | 22 | / | / | Immunofluorescence | R.F |
| 73 | Control | 32 | / | / | Immunofluorescence | R.F |
| 74 | Control | 22 | / | / | Immunofluorescence | R.F |

FCD = Focal cortical dysplasia; TLE = Temporal lobe epilepsy; PT = Peritumor cortical tissue of tumor induced epilepsy; Control = Control brain tissue with no history of epilepsy; Crypto = Cryptogenic; HS = Hippocampal sclerosis; CH = Cavernous hemangioma; GG = Ganglioglioma; PXA = Pleomorphic xanthoastrocytoma; PA = Pilocytic astrocytoma; PLNTY, Polymorphous low-grade neuroepithelial tumor of the young；TE = Traumatic epilepsy; R = Right; L = Left; T = Temporal lobe; F = Frontal lobe; P = Parietal lobe; OL= Occipital lobe; H = Hippocampus; * clinical data not recorded; / data not available.

**Supplemental Excel table titles and legends**

Supplementary table 2. Differentially expressed marker genes of patch-seq neuron clusters, related to Figure 1.

Supplementary table 3. Meta information for patch-seq neurons, related to Figure 1.
